# A Computational Model of Deep Brain Stimulation for Parkinson’s Disease Tremor

**DOI:** 10.1101/2024.04.26.591059

**Authors:** Sandeep Sathyanandan Nair, V. Srinivasa Chakravarthy

## Abstract

Parkinson’s Disease (PD) is a progressive neurological disorder that is typically characterized by a range of motor dysfunctions and its impact extends beyond physical abnormalities into emotional well-being and cognitive symptoms. The loss of dopaminergic neurons in the Substantia nigra pars compacta (SNc) leads to an array of dysfunctions in the functioning of Basal Ganglia (BG) circuitry that manifests into PD. While active research is being carried out in finding the root cause of SNc cell deaths, various therapeutic techniques are prevalent to manage the symptoms of PD. The most common approach in managing the symptoms is replenishing the lost dopamine in the form of taking dopaminergic medications such as Levodopa amidst its long-term complications. Another commonly used intervention for PD is deep brain stimulation (DBS), which is a invasive technique where an electrode is surgically inserted into the skull and a high frequency current of appropriate characteristics is delivered to the brain region. DBS is most commonly used when levodopa medication efficacy reduces and also in combination with levodopa medication that will help reducing the required dosage of medication prolonging the therapeutic effect. DBS is also a go to option when motor complications such as dyskinesias emerge as a side effect of medication. Several studies have also reported that though DBS is found to be effective in suppressing severe motor symptoms such as tremor and rigidity, it has adverse effect on cognitive capabilities. Henceforth it is important to understand the exact mechanism of DBS in alleviating the motor symptoms. A computational model of DBS stimulation for motor symptoms will offer great insights in understanding the mechanisms underlying the DBS and in this line in our current study we model a cortico-basal ganglia circuitry of arm reaching where we simulate healthy controls (HC) and PD symptoms as well as the DBS effect on the PD tremor. With DBS current characteristics of 220 pA, 130 Hz and 100 microseconds pulse-width we were able to see maximum therapeutic effect using our model. This model can be extended to accommodate cognitive dynamics in future so as to study the impact of DBS on cognitive symptoms and optimizing the parameters to get optimal performance effect across modalities.

## 1. INTRODUCTION

Parkinson’s disease (PD) is a severe neurodegenerative disease that affects a large percentage of the elderly human population and at present is second only to Alzheimer’s in terms of the number of people it affects [1], [2], [3]. With the progression of the disease, PD manifests into some of the cardinal symptoms such as tremor, rigidity, bradykinesia and even loss of balance ([3], [4], [5], [6], [7], [8]. The onset and progression of the disease has close link to the neuronal loss in substantia nigra pars compacta (SNc) [9], [10], [11] that unsettles the dopaminergic pathways in basal ganglia (BG). When contemplating symptom management strategies, the predominant approach often involves the replenishment of the dopamine loss, by the administration of dopaminergic medications such as levodopa (LDOPA) [12], [13]. Another commonly used technique is to stimulate the relevant brain region using an external current with appropriate characteristics, a technique known as Deep Brain Stimulation (DBS) [14], [15], [16]. Since prolonged usage of dopaminergic medications lead to motor complications like dyskinesias and the wearing-off effect, the number of people undergoing DBS surgery has increased significantly. Research suggests that DBS is useful not only in cases where the effectiveness of medication diminishes, or severe side effects of the medication occur, but also in reducing medication dosage [17]. Furthermore, when combined with LDOPA, it provides a highly effective therapeutic approach [18], [19].

DBS entails a procedure where an electrode is surgically implanted into the skull, delivering an electrical current with precise parameters into the subcortical region. While dopaminergic medications like Levodopa focus on rectifying dopamine deficiency within the BG pathways, DBS targets specific functional regions of the pallido-subthalamic circuitry and serves as a catalyst for exploration within the cortico-basal ganglia circuitry, essential for learning.

This exploration is particularly facilitated through the modulation of the pallido-subthalamic circuitry, which constitutes the indirect pathway of the basal ganglia. From a functional connectivity standpoint, cortical inputs reach the BG-Striatum’s input port, which then channels information to the output nucleus, GPi, through two pathways—one directly and another via the pallido-subthalamic circuitry. By applying DBS to the STN, an integral component of this circuitry, the intricate dynamics of information flow and modulation within the basal ganglia thalamocortical network are further elucidated.

Numerous studies have observed increased beta band oscillations and beta power in the STN region in PD patients. Experimental research has revealed that the interplay between cortical and basal ganglia (BG) structures, including the pallidum and subthalamic nucleus (STN), can induce beta rhythm oscillations throughout the cortico-basal ganglia system. These oscillations are commonly seen during various PD symptoms [20], [21], [22], [23]. The low dopamine levels lead to synchronous firings of both STN and GPe neurons [24], [25]. Theoretical studies have shown the relationship between the neuronal synchrony and collateral strengths in both STN and GPe neurons [26], [27]. DBS application mitigates synchronous neuronal communication and partially restores normal functioning of the indirect pathway circuitry, contributing to improved behavioral outcomes.

To delve into the dynamics of pallido-subthalamic circuitry, initial studies focused on studying the various firing characteristics by modulating the synaptic strength and connectivity patterns [28]. Later this single-compartment biophysical model expanded [28] by incorporating globus pallidus interna (GPi) and thalamus [29]. The relationship between PD tremor and the oscillations in pallido-subthalamic circuitry was discussed by various experimental studies [20], [30], [31], [32], [33] and also the relationship between the STN and cortical oscillations has been explored [34].

While the aforementioned models provide insights into neural activity and dynamics, it is crucial to comprehend how these effects translate into behavior, as the ultimate aim of DBS is the wellbeing of individuals with PD. Modeling behavior allows for a comprehensive assessment of the diverse impacts of PD and the potency of interventions like DBS. PD affects various behaviors, including motor functions such as tremors, rigidity, impaired coordination, and movement precision, as well as cognitive functions like decision-making and executive control. The impact of DBS on cognitive symptoms has been explored using a computational model of spiking neurons [35], [36] it was found that stimulation of STN worsened the decision-making performance in tasks such as Iowa Gambling task.

When it comes to motor tasks, modeling arm-reaching tasks has a particular significance. Reaching tasks are pivotal in daily activities and are intricately linked with motor symptoms experienced by PD patients. By simulating these tasks, researchers can gain insights into the specific motor deficits present in PD and evaluate the effectiveness of interventions like DBS in restoring motor function and improving overall motor performance.

In the current study we focus on modelling the cortico-basal ganglia circuitry to replicate arm-reaching behavior and investigate the therapeutic efficacy of DBS on such behavior. Earliest efforts in modelling coordinated reaching movements started with control-system based loops [37], [38], [39], [40], [41][42]. While this research didn’t focus on the underlying neural mechanisms, parallel research were conducted on neural substrates and neural mechanisms involved in reaching [43], [44], [45]. Soon reinforcement learning-based models were in use [46] and in our study this arm model [46] was integrated into a basal ganglia thalamocortical model consisting of the oscillatory model of STN-GPe network [47] to simulate motor movements at the behavioral level [48], [49]. In order to accommodate the DBS effect on performance, we replace the rate coded model of pallido-subthalamic circuitry with a spiking neuron model and study DBS effects.

This outline of the paper is as follows. Section 2 discusses the materials and methods followed by Section 3 that describes the results followed by the discussion section. In the methods section we will describe the cortico-basal ganglia model and its key components. We will then describe the mechanism to simulate PD condition followed by the DBS intervention. After the methods section, we highlight some of our important results from the model that is then followed by the discussion.

## 2. MATERIALS AND METHODS

We use a cortico-BG (CBG) model that controls a two-linked arm model in order to simulate reaching movements as shown in Fig.1. In this section we will first introduce the CBG model, and DBS intervention. We will describe the arm model in the supplementary section.

### 2.1. Cortico-basal Ganglia Model

The CBG model used in our study comprises of an outer sensory motor loop, which interacts with the BG circuitry. The sensory motor loop comprises of an arm model, proprioceptive cortex (PC), motor cortex (MC), the prefrontal cortex (PFC) and the BG. The kinematic arm model performs reaching movements based on the activations it receives and the PC estimates the current arm position and sends the feedback to MC, which integrates this signal along with the goal information from the PFC and the error corrected signal from the BG, which does its own internal processing before sending back the corrected signal to the motor cortex. The motor cortex then sends the next motor command to the arm via the spinal motor neurons and this process continues until the arm reaches the target or the time out is reached. More details about the sensory motor loop and the arm model is discussed in the supplementary material. The MC acts as the layer of intersection between the outer sensory motor loop and the BG circuitry. The communication between the MC and the BG is crucial in exploring and understanding the PD dynamics.

The BG consists of the Striatum, which is the input nucleus, Globus Pallidus interna (GPi), which is the output nucleus and the GPe and STN that constitutes the indirect pathway.

The Subthalamic nucleus (STN) and Globus pallidus interna (GPi) that constitute the indirect pathway is modelled using the spiking neuron model (Izhikevich), while the rest of the BG nuclei is modelled using the rate coded model.

#### 2.2.1 The MC-BG interaction

Effective interaction between BG and cortex plays a crucial role in motor acquisition and performance. This interplay facilitates learning, leading to decision-making scenarios where competing signals—one facilitating and the other inhibiting—are evaluated. This evaluation forms the basis of selecting the best action and is facilitated by GPi mechanism [50], [51]], [52], where inputs via two parallel pathways – the D1 and D2 combine. The flow of information through the D1 and the D2 pathways are modulated by the dopamine signal from the SNc [53]. Depletion of dopamine signals due to neuronal death results in motor impairments, which are then managed using dopaminergic medication or DBS.

The MC receives inputs from the PFC, the PC and the BG and the total input received at the MC, *I*^*MC*^, is as given in equation <u>(1)</u> below. The dynamics of PC and PFC is given in the supplementary section.

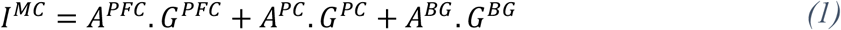

The dynamics of the MC neurons is defined by the continuous attractor neural network (CANN) and the MC output (*G*^*MC*^) is as given in equation (2) below,

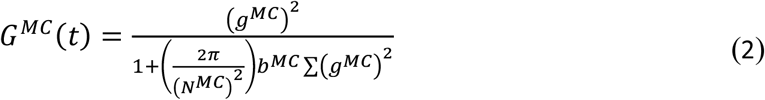

where, *N*^*MC*^ defines the network size of MC, *b*^*MC*^ is a constant and *g*^*MC*^ represents the intrinsic state of the nodes in MC and is as given in the equation (3) below,

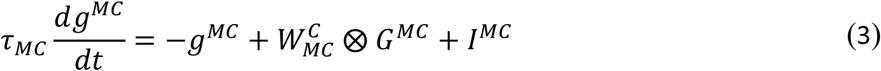

where, the weight kernel 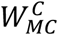 represents the lateral connectivity among the MC neurons and *⨂* represents the convolutional operation.

The MC input is presented to the Striatum (STR), which routes the signal to the GPi directly as well as through the STN-GPe network. The signal projecting from the D1 medium spiny neurons (MSNs) of the Striatum to GPi is *y*^*D1*^ as given in equation <u>(4)</u>, whereas signal projecting from the D2 MSNs of the Striatum to GPe is given as *y*^*D2*^ in eqn. <u>(5)</u>. The *D1*_*λ*_ and *D2*_*λ*_ in equations (6) and (7) represent the sigmoidal activation of the D1 and D2 striatal neurons.

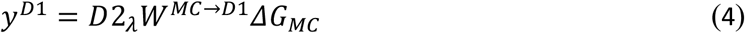

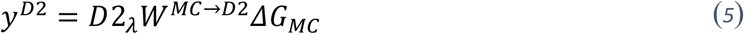

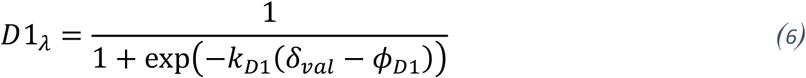

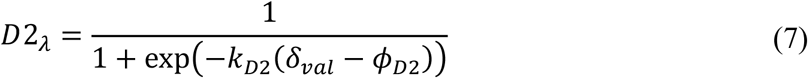

where the weights *W*^*MC→D1*^ and *W*^*MC→D2*^ represent the weights between MC and STR, *ΔG*_*MC*_ is the input received by STR from MC, *δ*_*val*_ is the value difference that modulates the BG pathways, *k*_*D1*_ and *k*_*D2*_ are the sigmoidal gains, where *k*_*D1*_ *= −k*_*D2*_ and *ϕ*_*D1*_ and *ϕ*_*D2*_ are the thresholds used in the sigmoidal function for *D1*_*λ*_ and *D2*_*λ*_, respectively.

The quantity *δ*_*val*_ used here is termed the *value difference* and it is computed as given in equation (8) below,

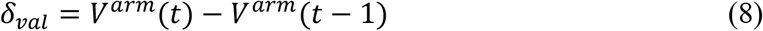

*V*^*arm*^(*t*) mentioned in eqn. (8) represents the value of the current position of arm at time ‘t’ and is obtained by probabilistic gradient ascent over the value function (Chakravarthy & Moustafa, 2018a; Muralidharan et al., 2018; Nair et al., 2022) performed by BG as given in equation (9) below. *X*^*arm*^ and *X*^*targ*^ are the current arm position obtained from PC and the target goal position obtained from PFC respectively. The value function, *V*^*arm*^(*t*) is obtained as given in equation (9) below.

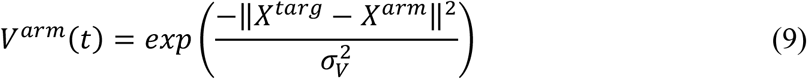

where,, the spatial distance within which the value function demonstrates sensitivity for that particular target is given by *σ*_*V*_.

#### 2.2.2 The STN-GPe subsystem

The D2 MSNs of the STR project to the GPe, and the current received at the GPe neurons from the STR is as given in equation (10) below.

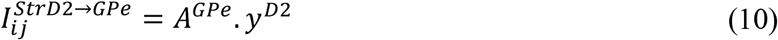

With *A*^*Gpe*^ representing the weight between the STR and GPe. The total incoming current at the GPe is as given in equation (11) below.

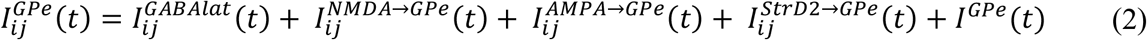

The GPe neurons also receives inputs from its own neighbouring neurons, which is as given in eqn. (12) below and the corresponding neurons in STN, as given in equations (13 and 14) below.

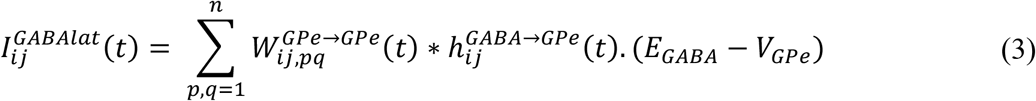

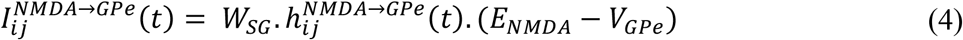

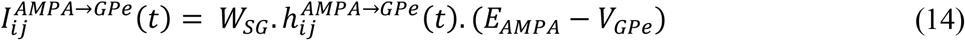

In the above equations *V*_*Gpe*_ represents the voltage across the membrane for the GPe neurons as described in equations (15-17) below. The STN-GPe network consists of 2D arrays of spiking neurons modelled using Izhikevich equations. When the membrane voltage reaches *V*^*peak*^, the variables are reset as shown below.

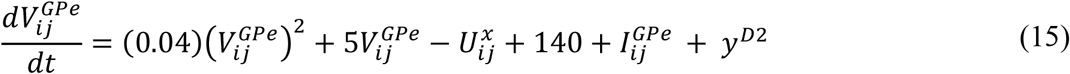

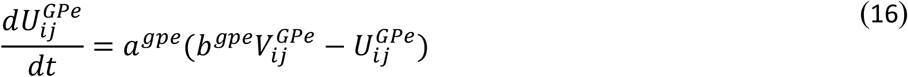

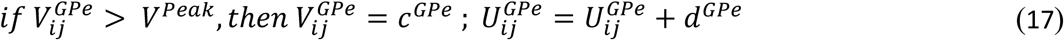

Also, 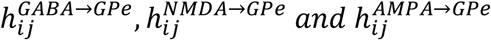 represent the gating variables as shown in equations (18-20) below.

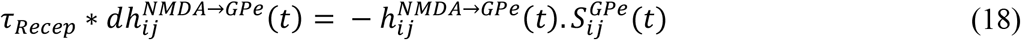

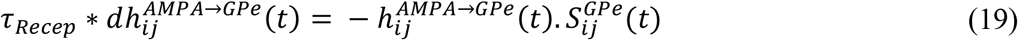

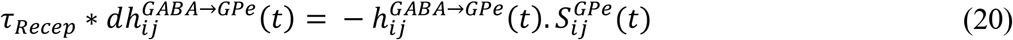

*I*^*GS*^ is the current that goes from the GPe neurons to the corresponding STN neurons that is as shown in equation (21) below. Here *W*_*GS*_ denotes the weights between STN and GPe neurons, 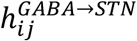 regulates the gating between GABA and STN, *V*^*STN*^ is the voltage across the membrane for the STN neurons and *E*_*GABA*_ is the voltage across the membrane at rest for the GPe neurons.

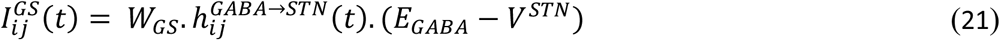

Along with *I*^*GS*^, the lateral currents from the neighbouring STN neurons constitute the total current received at each STN neurons, which is governed by equations (22-24) below.

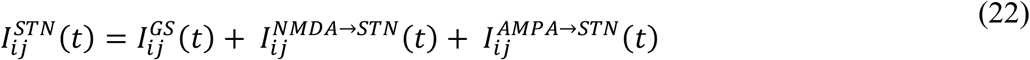

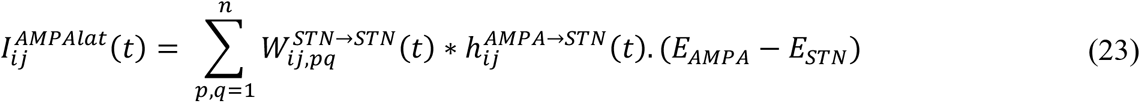

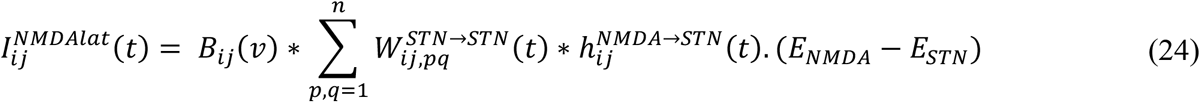

In the above equations, the terms *h*^*AMPA→STN*^, *h*^*NMDA→STN*^ and *h*^*GABA→STN*^ represent the gating variables and their dynamics are as shown in the following equations (25-27).

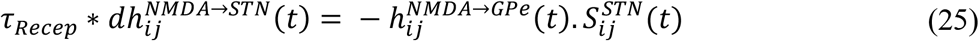

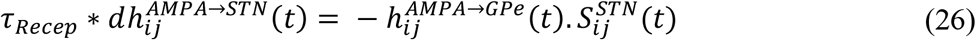

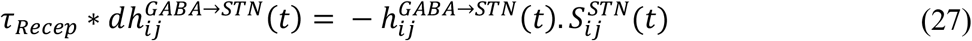

The membrane potentials of the STN neurons are described in equations (28-30) below. The terms 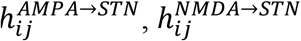 and 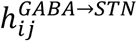 represent gating variables and their dynamics are as shown in the following equations.

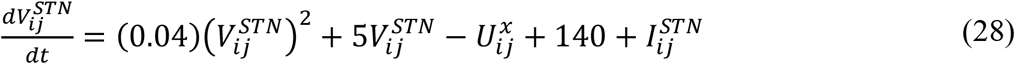

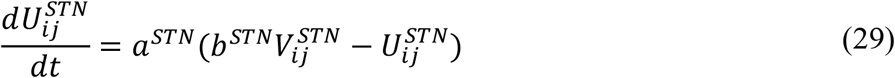

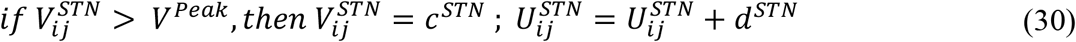

The local field potential (LFP) of the STN is calculated as shown in equation (31) below,

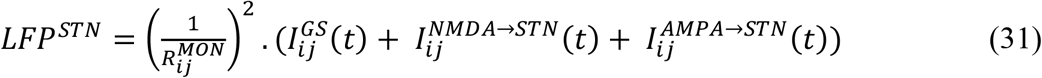

The term 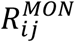 in the above equation is the distance between the (*i, j*)^*th*^ neuron and the recording point.

#### 2.2.3 Simulating the PD condition

To replicate the symptoms of PD, DA value is decreased from *DA*_*HC*_ to *DA*_*low*_. In other words, we prevent the *δ*_*val*_ from rising beyond *DA*_*low*_ as given in equation (32) below.

If PD=1 then,

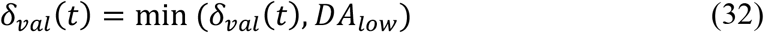

The DA value also influences the lateral connections of STN and GPe nucleus and also the interconnectivity (*W*_*SG*_ *and W*_*GS*_) between the corresponding neurons of STN and GPe as shown in equations (33-37) below.

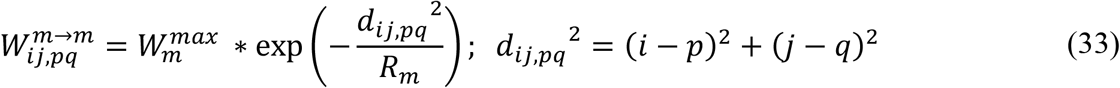

where m in the above equation represents STN/GPe. 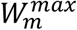 is the maximum connectivity strength among the neurons, *R*_*m*_ defines the radius of the neighbourhood, d is the distance between two neurons in the subpopulation and (i,j,p,q) represent the indices of the neurons.

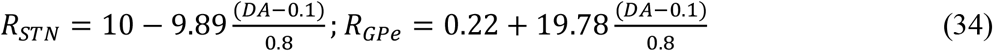

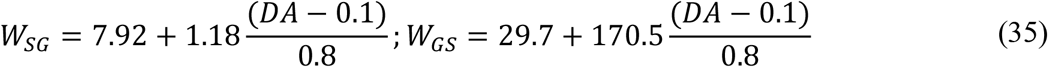

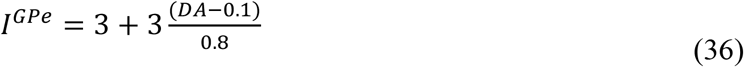

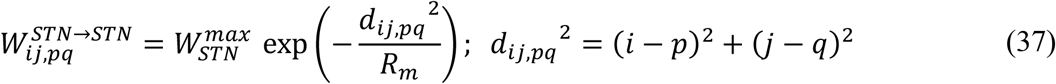

##### 2.2.3.1 Simulating Tremor and Rigidity conditions

For simulating the tremor and rigidity conditions, we modulate the connectivity strength between STN and GPi using the gain factor *A*^*D2*^, mentioned in equation 38. For tremor, a relatively higher value (2) of *A*^*D2*^is chosen, whereas for rigidity a lower value (<0.4) of *A*^*D2*^is chosen.

#### 2.2.4 The STN to GPi connection

Inputs from STN to GPi are taken after converting the spike data of STN neurons into rate codes. The mean rate of firing of the STN neurons is calculated as shown in equation (38) below.

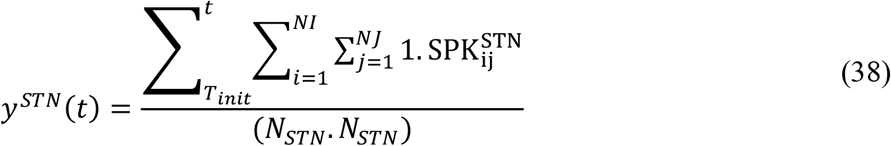

where *y*^*STN*^ *is the* average firing rate of the STN neurons for a simulation time of 1sec, 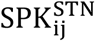 is the spike data of neuron at location (i, j) in the network, N is the total number of neurons (NI*NJ), T=simulation time (1sec). *T*_*init*_ *= t − WS* if t is greater than or equal to WS, else *T*_*init*_ *= t* .

#### 2.2.4 The BG to MC connection

The input through the direct projections from the D1 MSNs of the STR (*Y*^*D1*^) and the output of STN (*y*^*STN*^) are combined at GPi (*y*^*GPi*^) as shown in equation (39) below before forwarding to the thalamus.

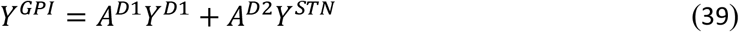

The dynamics of the thalamic neurons is modeled as a CANN and the thalamic output is as given by equation (40) below,

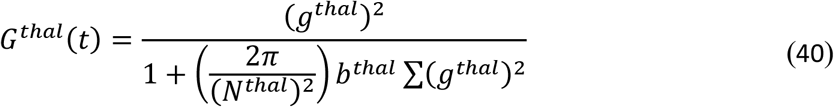

where, *N*_*thal*_ defines the thalamic network size, *b*_*thal*_ is a constant, *A*^*D1*^ *and A*^*D2*^ are the respective gains associated with the two pathways, and *g*_*thal*_ represents the intrinsic state of thalamic neurons as given by equation (41), .

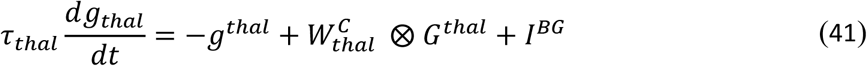

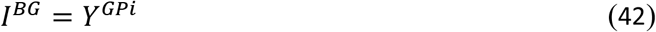

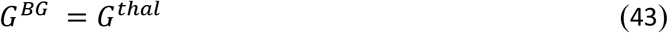

where, the weight kernel, 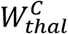, represents the lateral connectivity strength among thalamic neurons, *I*^*BG*^ is the input from GPi to thalamus coming from BG, *Y*^*GPi*^ and *G*^*thal*^ are outputs of GPi and thalamus respectively and *G*^*BG*^ is the thalamic output that goes to the MC.

#### 2.2.3 Parameter selection

The list of parameters used in the model and their corresponding values along with their description is given in the supplementary section. Also, the learning mechanisms are described in detail in the supplementary section.

#### 2.2.4 DBS effect

There are various targets used for DBS and among them GPi and ventralis intermedialis (Vim) of thalamus are commonly used apart from STN [56], [57], [58]. However, STN DBS is most commonly used for PD [16] considering better therapeutic effects.

The pulsatile current of appropriate parameters (amplitude, frequency and pulse duration) mimicking the clinically delivered DBS [59]effect is simulated in our model. The current is applied to the centre most neuron (position in the 2D lattice (*i*_*m*_, *j*_*m*_) and the spread of the current to the neighbouring neurons is modulated by a gaussian distribution [60], [61] with variance (*σ*_*DBS*_) as shown in equation (44) below.

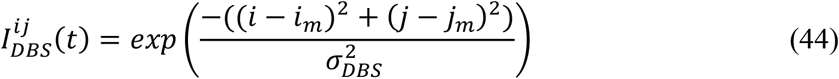

Where 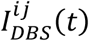 is the effective DBS current received by the neuron at location (i,j).

## 3. RESULTS

Here, we showcase the performance of the model by simulating both the healthy (HC) and PD condition and observe their effects on arm reaching task (Figure 4, 5). In order to simulate the HC and PD conditions we modulate the dopamine dependent control parameters in our cortico-basal ganglia model neuron model, especially the amount of current flowing from the D1 and D2 striatal neurons and also the parameters in the STN-GPe subsystem as detailed in the section below.

### Parameters controlling the firing patterns and Synchrony in STN-GPe

The neuronal firings and synchrony of the STN and GPe populations of neurons are tuned using various parameters such as the lateral connection spread of the GPe neurons(*R*_*Gpe*_), the lateral connection spread of the STN neurons (*R*_*STN*_), the spread of laterals in STN and GPe neurons, the interconnections between STN and GPe (*W*_*SG*_ *and W*_*GS*_, the dopamine availability, the striatum to GPe current(*I*^*Gpe*^) and the STN current. Dopamine signal (DA) modulates the lateral connectivity of both the STN and GPe populations as well as the interconnections between the two neuronal population as well as the input current to GPe as shown in equations (34-37).

### Neuronal Firings and Synchrony during Healthy and PD conditions

The variations in firing rate and synchrony with respect to the dopamine levels are as shown in Fig. 2 below. As the dopamine level increases from 0.1 to 0.9, the firing rates of the STN neurons decreases, whereas the firing rates of the GPe neurons increases (Fig. 2a). Also, the synchrony among the neurons of both STN and GPe decreases with increasing dopamine levels (Fig. 2b).

**Figure 1.**
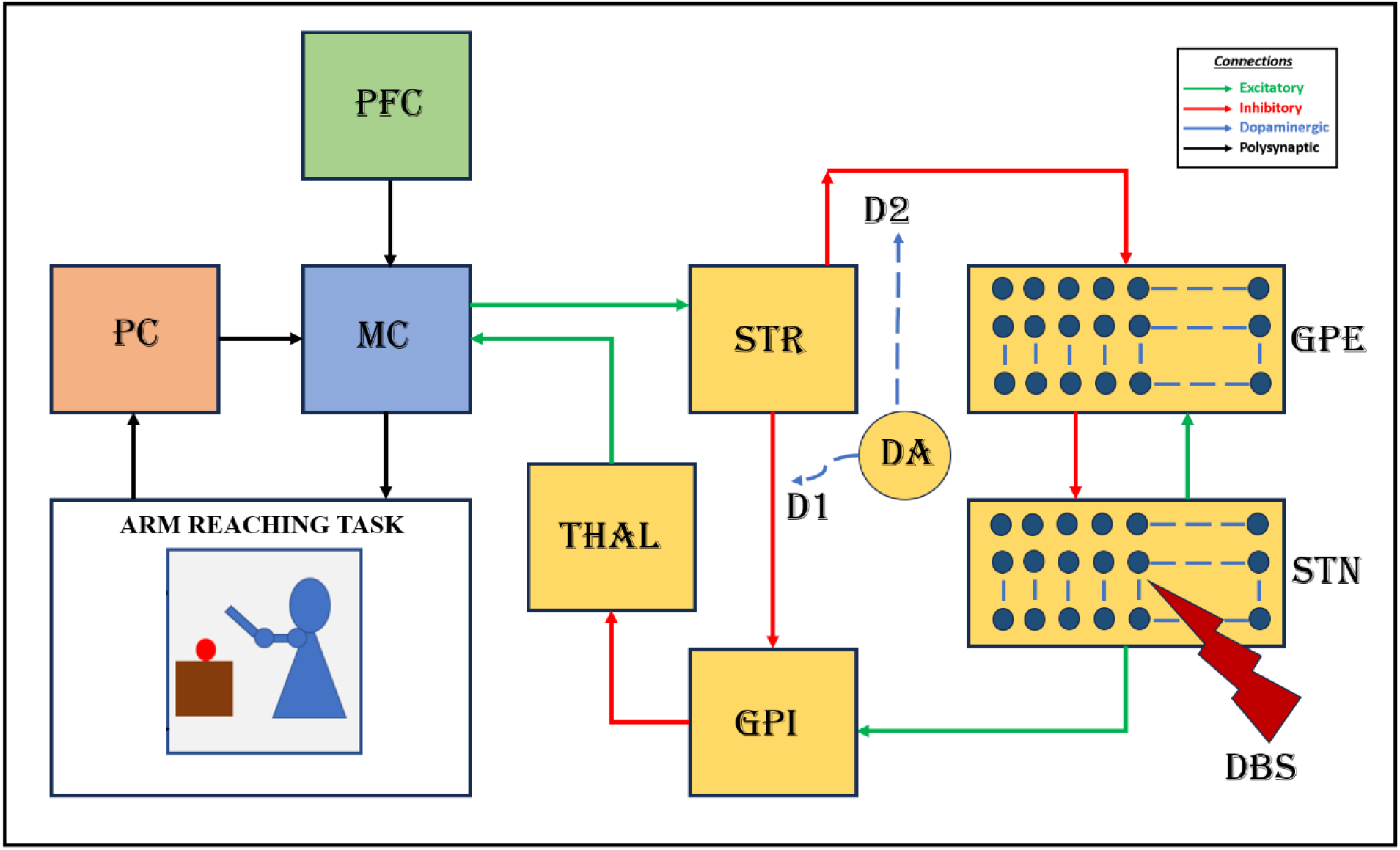
Block diagram of the proposed cortico-basal ganglia model. The model consists of a 2-link arm model, the proprioceptive cortex (PC), the prefrontal cortex (PFC), the motor cortex (MC), and Basal ganglia (BG). Here the input nucleus Striatum, the output nucleus Globus pallidus interna (GPi), the Globus pallidus externa (GPe), the subthalamic nucleus (STN), and the thalamus (THAL) constitute the BG. MC integrates the inputs received from the prefrontal cortex (PFC) and the proprioceptive cortex (PC) along with the feedback signal from BG and sends the signal to the arm via the spinal motor neurons.

**Figure 2.**
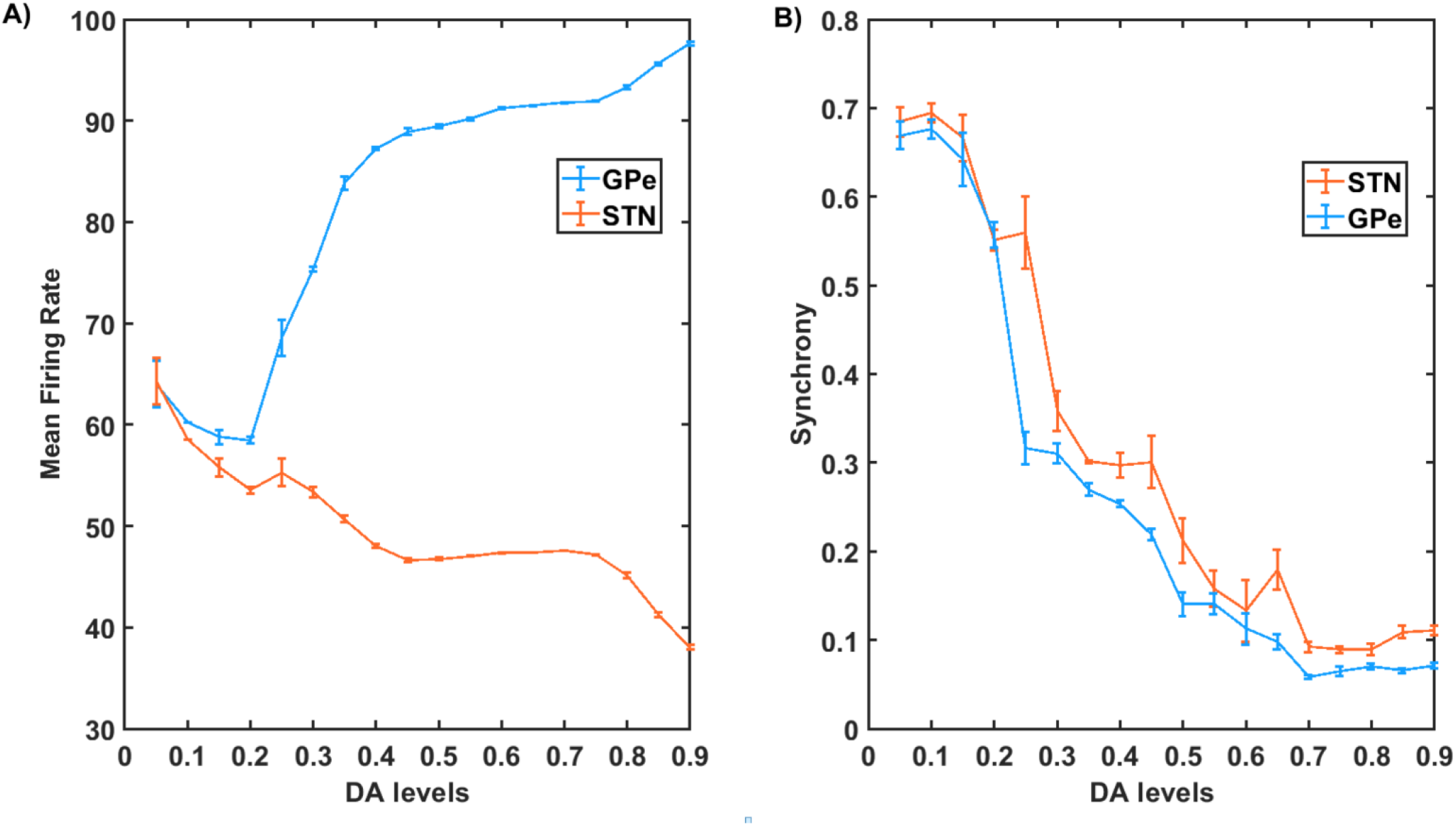
A) The firing rates of STN and GPe neurons for various values of DA levels are shown. The blue line represents the STN and the orange line represents the GPe neurons. B) Synchrony within STN and GPe nuclei. Again, blue and orange lines represent the GPe and STN neurons respectively. Synchrony keeps decreasing with increasing DA levels. The mean and variance values for the above plots are calculated over 5 epochs.

We select two DA values 0.1 and 0.9 to simulate PD and HC conditions. The neuronal firings and synchrony of the STN and GPe populations of neurons under both these conditions are as given in the Fig. 3 below. Fig. 3(a-f) represents the dynamics under the healthy condition, whereas Fig. 3(g-l) represents the dynamics of the PD condition. Under the healthy condition (HC) both STN and GPE neurons exhibit regular spiking as seen in Fig. 3aand Fig. 3dand the activities of the neuronal subpopulation in both STN and GPe exhibits asynchronous firings as seen in Fig. 3bandFig. 3e Fig. 3cand Fig. 3fshows the synchrony among the STN and GPe neuronal population as a function of time. During PD condition the dopaminergic neurons in the SNc die and in our model, we simulate the same by reducing the dopamine level from 0.9 to 0.1. The reduction in dopamine level influences the STN-GPe circuitry via the lateral connections of STN and GPe and the interconnections between the STN and GPe. Fig. 3gandFig. 3j shows that due to this influence, the STN and GPe fires in bursty mode and there is an increased synchrony among both the STN and GPe neurons as shown in Fig. 3handFig. 3k. Fig. 3(c, f, i, l) represents the synchrony among the neurons of the respective neuronal populations.

**Figure 3.**
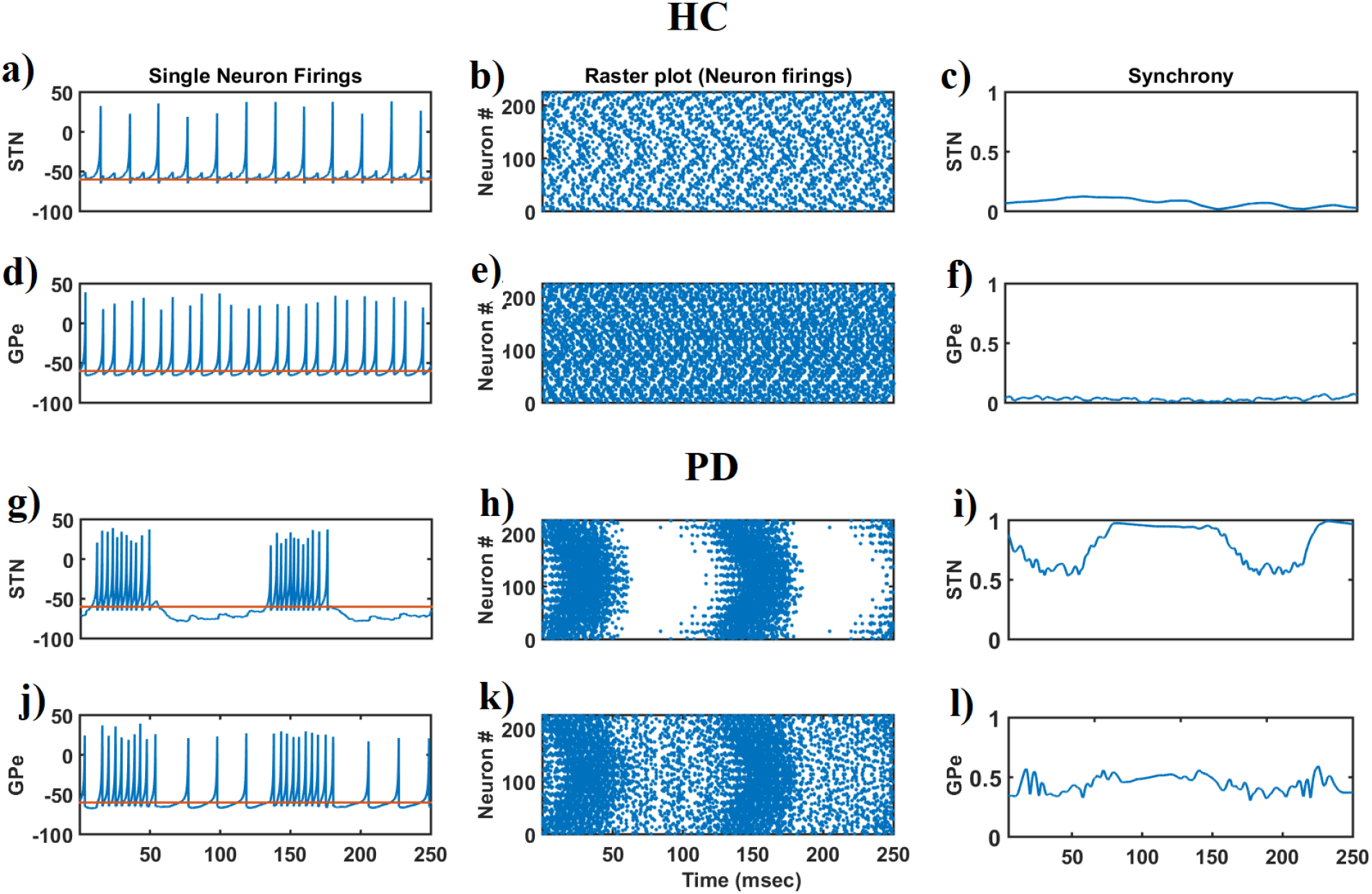
The firing of a single STN and GPe neuron under HC is shown in 3a & 3d. Under HC condition both the STN and GPe neurons exhibit regular firings. The firing of a single STN and GPe neuron under PD condition is shown in 3g & 3j. 3b & 3e shows the raster plot of the STN and 3h & 3k shows that of the GPe neurons under PD conditions. 3c & 3f indicates the synchrony of STN and GPe neurons under healthy conditions whereas 3i & 3l indicates the synchrony of STN and GPe neurons under PD conditions.

### Arm reaching performance during Healthy and PD conditions

Reaching movements are simulated using the model described in Fig. 1with DA level set to 0.9. The blue line in the Fig. 4represents the reaching performance of the HC. It can be seen that during a normal condition (HC) the arm reaches the target in a consistent manner with the velocity of the arm forming a bell curve Fig. 4cwith respect to time where the speed increases initially until it reaches a peak and then slowly reduces as the arm approaches the target. The distance to target slowly reduces, resembling a waterfall curve, as it reaches the target (Fig. 4b). Acceleration of the arm during the reaching task is as shown in (Fig. 4a), where we do not see any significant peaks in the tremor frequency (4-10 Hz) band in the spectrum of the arm acceleration, while we do see a significant peak around 7 Hz in case of a PD tremor condition represented by the orange line in (Fig. 4a). Under PD tremor condition we can also see that the arm never reaches the target and it keeps fluctuating as shown in the orange curve (Fig. 4B). Also, the velocity of the arm keeps increasing and decreasing while the tremor is experienced as shown in the orange curve (Fig. 4C)

**Figure 4.**
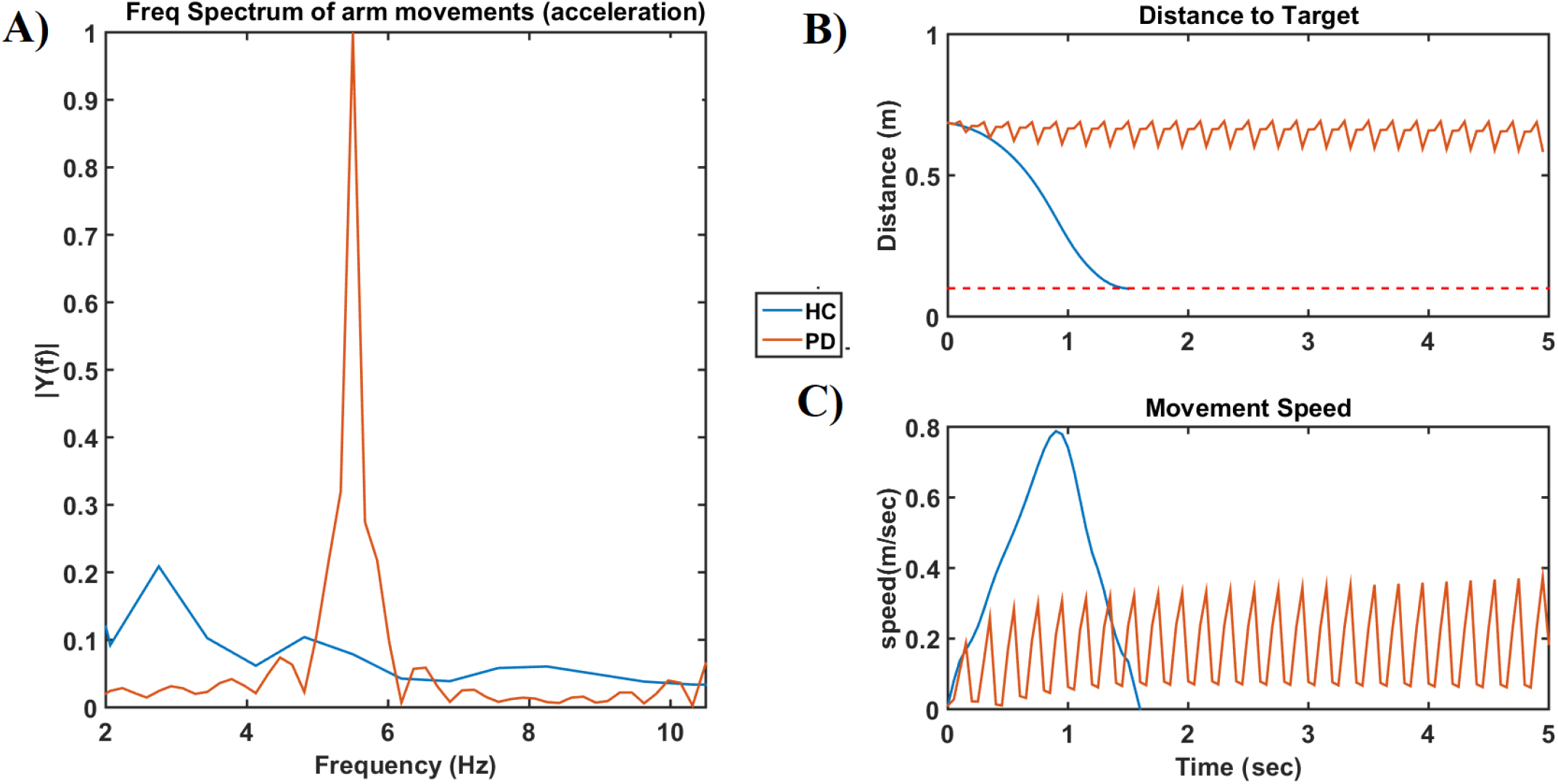
A) The frequency spectrum of the acceleration of the arm movements is given. The blue line represents the healthy controls (HC) and the orange line represents the PD tremor condition in all (a, b, c). B) This plot shows the distance to the target as the time progresses. C) The velocity of the arm movement where the curve follows a bell curve under HC and keep oscillating under PD tremor condition.

The trajectory of the arm is as shown in the Fig. 5below. During a normal (HC) the arm reaches the target in a consistent fashion as indicated by the blue arm, whereas during a PD tremor condition the arm keeps fluctuating as shown by the yellow arm and during a rigidity case the arm hardly moves from the starting position as shown by the green arm. The red dotted circle around the target position is the region where the arm is considered to have reached the target within its area.

**Figure 5.**
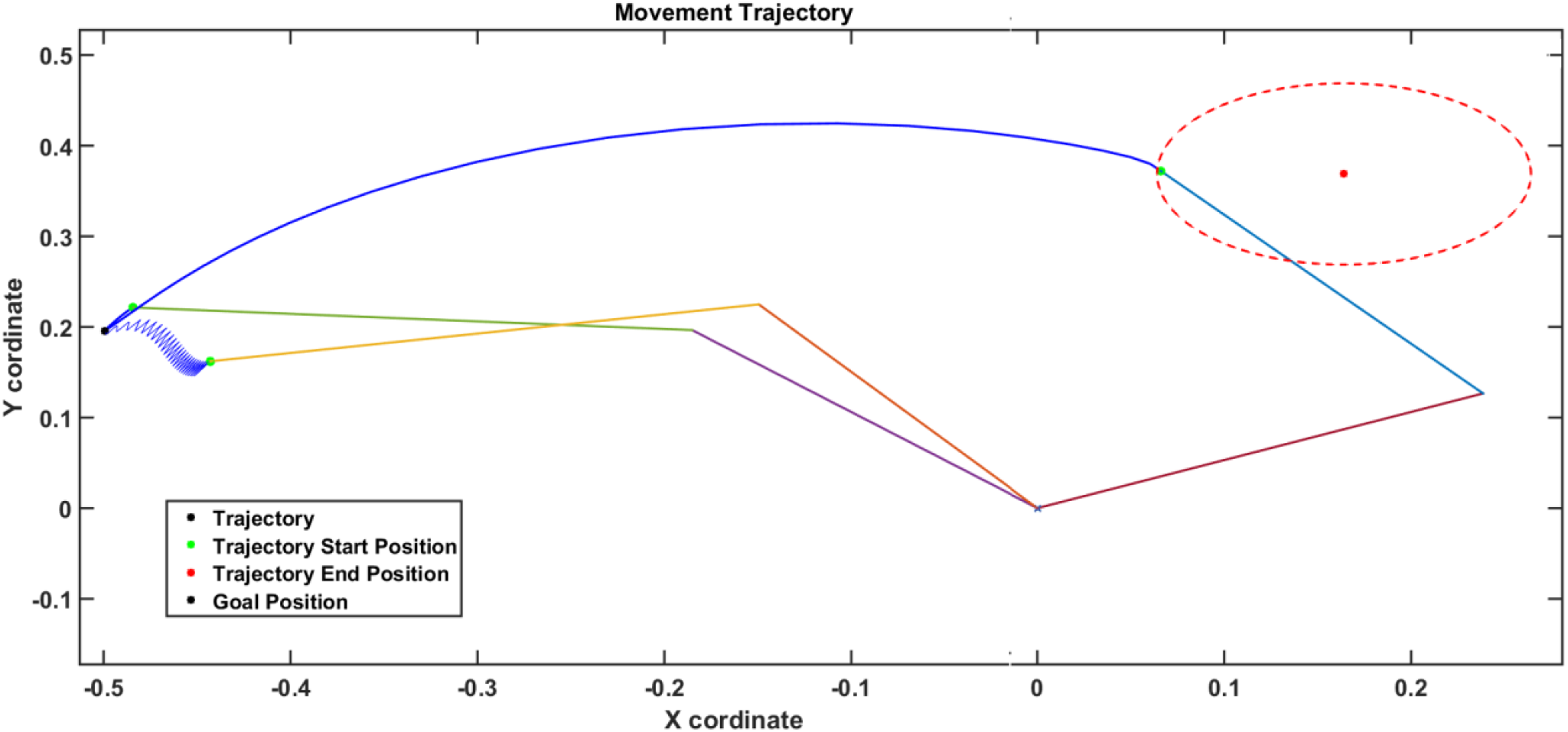
The trajectory of the arm movements is given. The blue line (for arm) represents the healthy controls (HC) and the yellow line (for the arm) represents the tremor condition and green line (for the arm) represents the rigidity condition. In case of HC the reaching is successful, whereas in case of tremor and rigidity it is not.

Under HC the distance to target of the arm steadily decreases and reaches the target (∼1.5 secs) as shown in the green line in (Fig. 6a), while under tremor condition we can also see that the arm never reaches the target and it keeps fluctuating as shown in the orange curve. In case of rigidity the arm hardly moves and distance to target is a steady flat line as shown by blue cure. (Fig. 6b) shows the velocity of the arm while performing the reaching, and as seen the green line representing HC resembles a bell curve, where the velocity increases until it peaks and then gradually decreases. Also, the velocity of the arm keeps increasing and decreasing under tremor condition (orange line) and hardly raises under rigidity condition (blue line).

**Figure 6.**
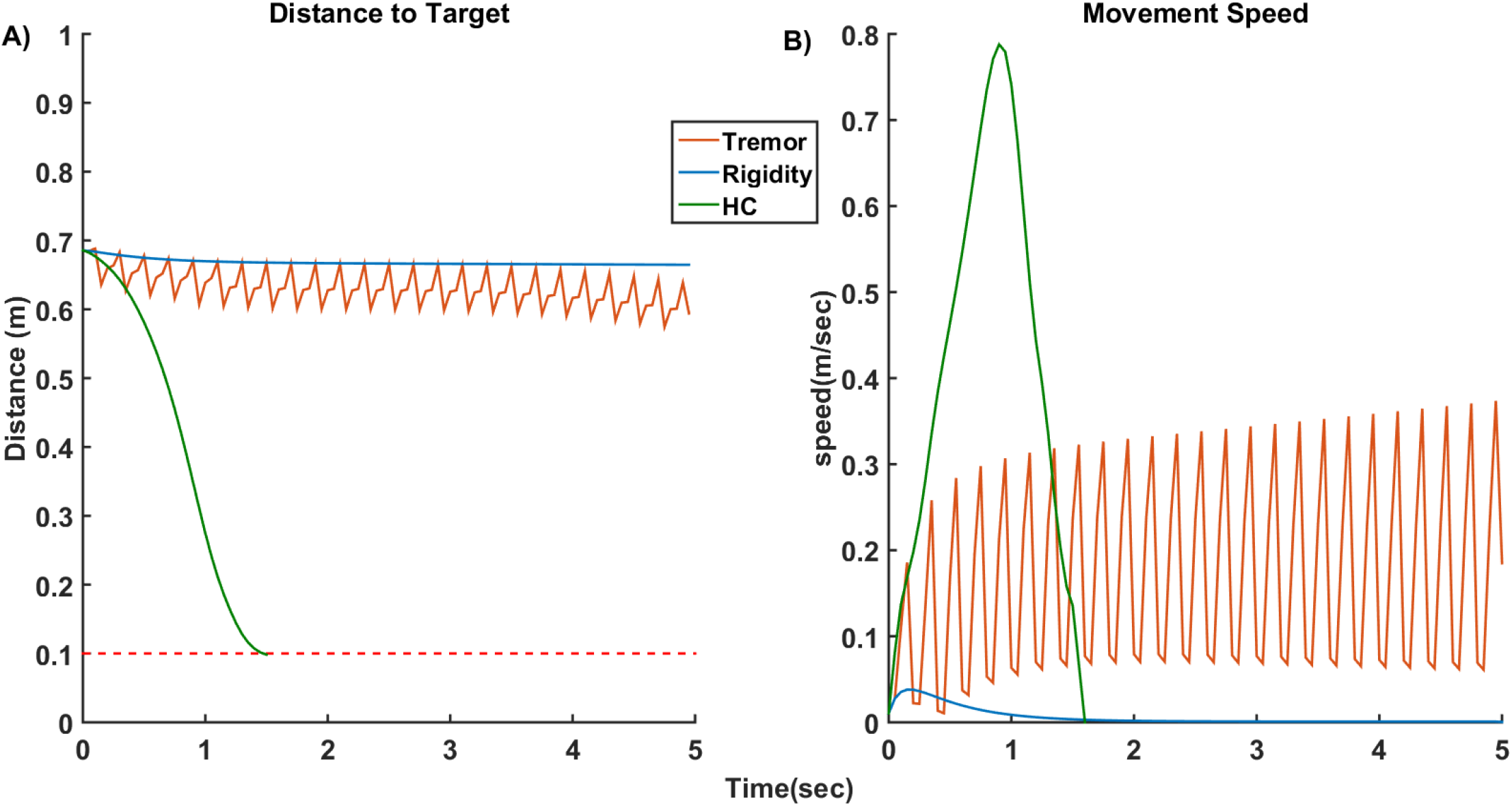
A) This plot shows the distance to the target as the time progresses.. B) The velocity of the arm movement where the curve follows a bell curve under HC and keep oscillating under PD tremor condition. And hardly rises and quickly dies down under Rigidity condition. The green line represents the healthy controls (HC), the blue line represents the rigidity condition and the orange line represents the tremor condition

During the tremor condition the neuronal population of the STN subsystem fires in a synchronous manner and hence the local field potential (LFP) of the STN is highly periodic and higher in amplitude (green line) as shown in Fig. 7, whereas the amplitude of the LFP signal of HC and DBS treated conditions represented by violet and blue lines is comparatively much smaller. Looking closely at the dynamics of STN-GPe, the frequency spectrum of the LFP of the STN neuron population reveal that there is a significantly higher power observed in the beta frequency band (13-35 Hz) as shown in Fig. 7C. During the tremor condition, the beta peaks are also accompanied by another peak at the theta band (4-11 Hz). This is inline with the observations in the experimental study [62].

**Figure 7.**
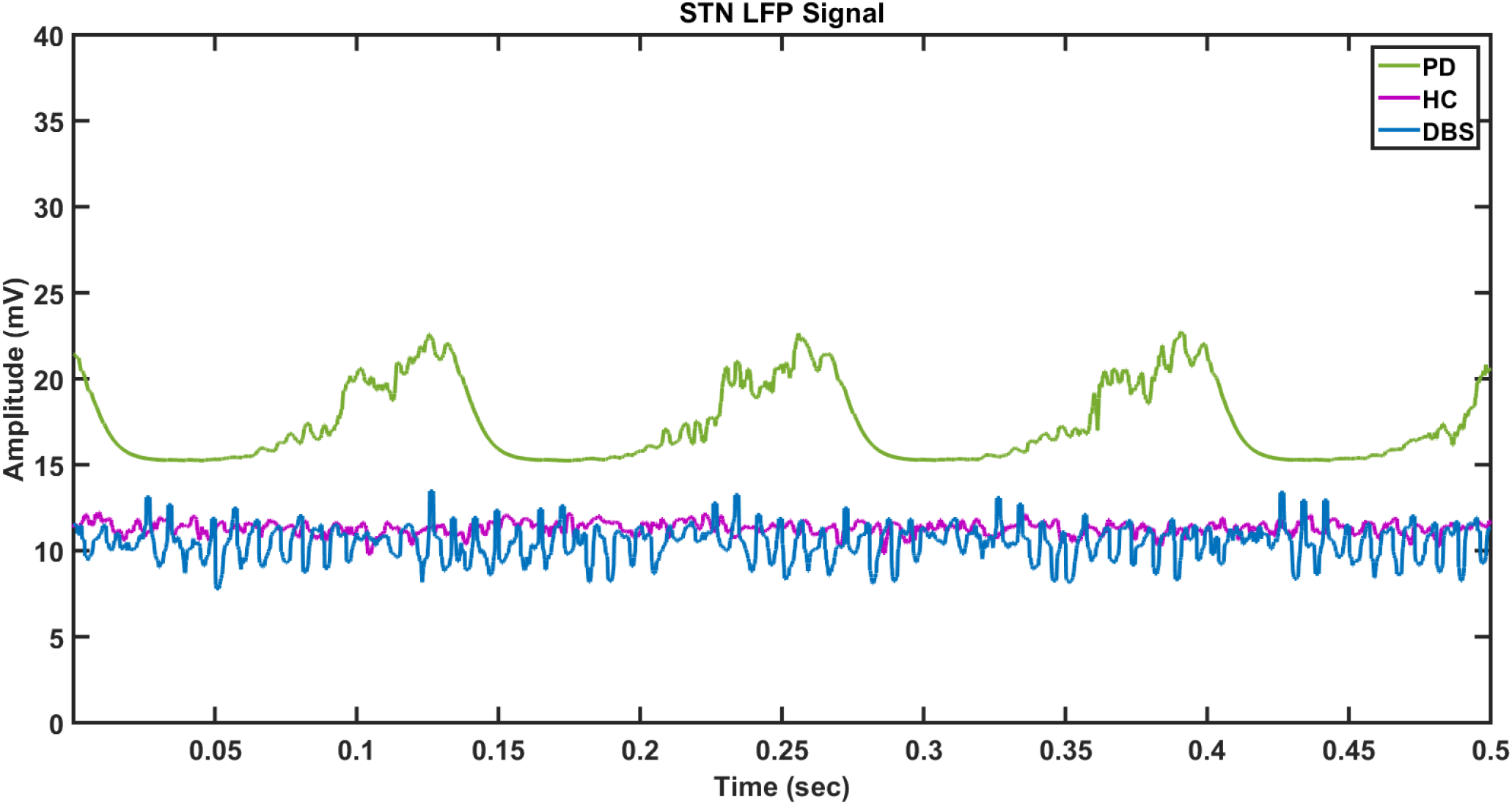
The local field potential of STN neuronal population is shown. The violet curve indicates the HC, the green line indicates the PD condition and the blue line represents the DBS treated condition.

**Figure 7.**
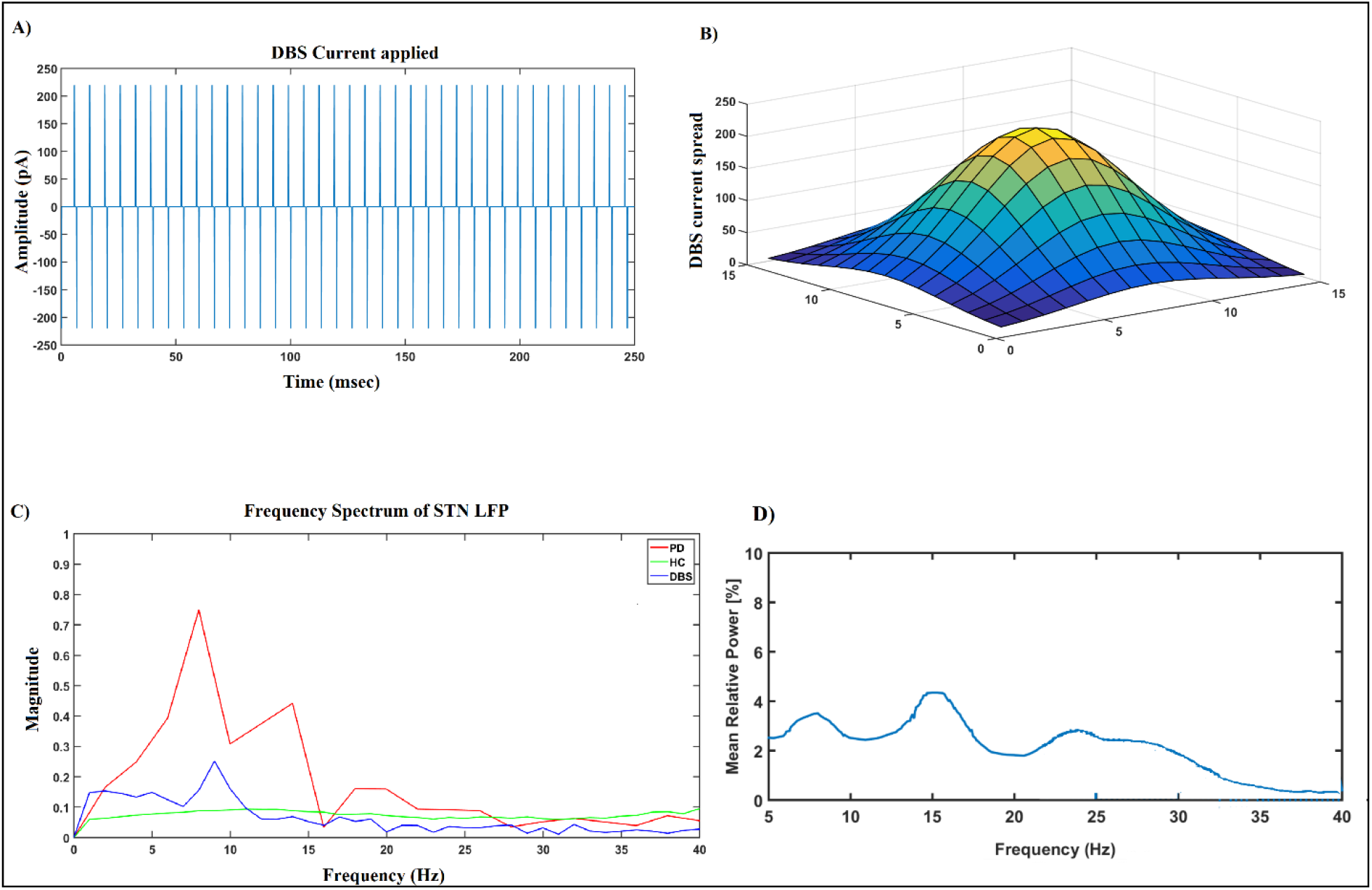
A) DBS current applied to the centre most neuron in the STN population. B) The spread of the current in nearby neurons. C) The FFT of the local field potential of the STN population. D) The mean relative power of the LFP of the STN redrawn as recorded in the experimental studies (Kuhn et al., 2008) [62].

### Effect of Deep brain stimulation (DBS)

As it is evident that significant beta peaks in the LFP of STN is a signature of PD, attempts have been made to suppress this beta peak. The DBS facilitates suppression of beta peaks by injecting a high frequency current of appropriate amplitude and pulse duration. The DBS current used in our simulation is as shown in Fig. 8A. A biphasic pulsatile current of 130 Hz, 220 pA and 100 microseconds pulse duration is applied to the centre most neuron of the STN sub population and the effect of the DBS current on the neighbouring neurons is modeled as a gaussian spread as shown in Fig. 8B. In our study we check the impact of STN DBS on reaching performance under tremor condition. The current is applied to the centre most neuron. We use biphasic, single contact stimulation.

**Figure 8.**
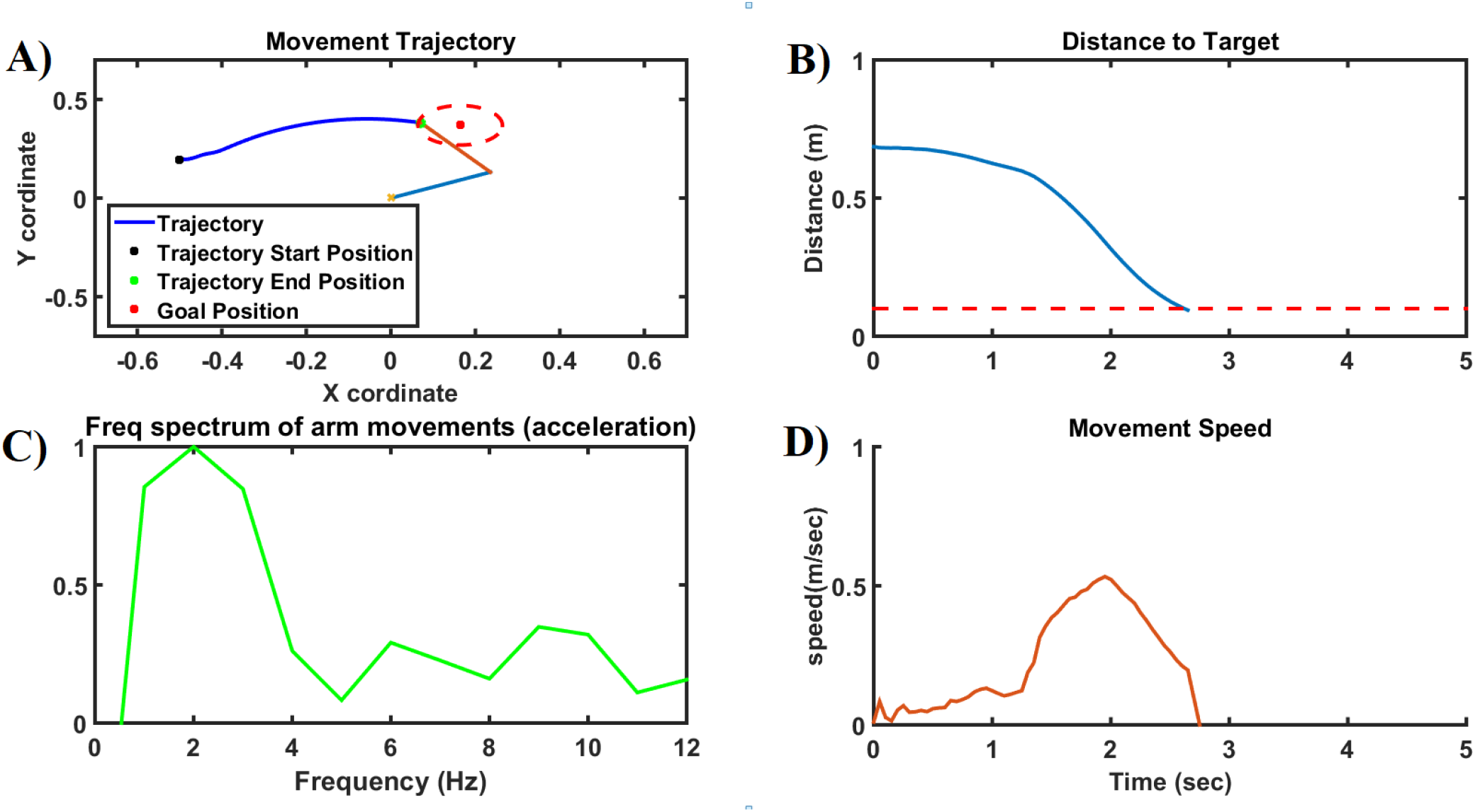
The trajectory of the arm movements is given. The blue line (for arm) represents the healthy controls (HC) and the yellow line (for the arm) represents the tremor condition and green line (for the arm) represents the rigidity condition. In case of HC the arm reaches the target, whereas in case of tremor and rigidity the arm doesn’t reach the target.

### Effect of DBS on reaching performance

The DBS stimulation restores the reaching performance of the arm movement as shown in 9 below. We can also observe that compared to the HC performance in Fig. 4 and Fig. 5, the performance still shows a vast improvement and the arm was able to reach the target with slightly more time (2.6 sec) compared to the HC (1.6 sec). Fig. 9A shows the movement trajectory for a DBS treated condition and it can be seen that the arm reaches the target after a period of time. Fig. 9B shows the distance to the target as a function of time and we can see that the distance to the target keeps reducing with time. Fig. 9Cshows the frequency spectrum of the acceleration of the arm movements during the reaching task and we can see that the power in the region between 4-10 Hz band is significantly reduced. Fig. 9Dshows the velocity curve during reaching movement and it can be seen that it takes a while to attain the peak velocity followed by a gradual reduction until the arm reaches the target.

## 4. DISCUSSION

The focus of this study is to develop a cortico-basal ganglia model, aiming to simulate the DBS therapeutic effects on PD motor symptoms, most importantly PD tremor. Since PD tremor was found to depend sensitively on the synchronized firing dynamics of the STN-GPe neurons [63], [64], [65], [66], [67]and DBS action that aims to suppress tremor is expected to supress the synchronized peak in STN-GPe, in the current model, we use a spiking neuron model of STN-GPe system.

The cortico-basal ganglia model used in this study, is based on concepts from Reinforcement Learning and is based some of our earlier work [68].

Central to our model is the idea that the cortico-basal ganglia system achieves movement control by performing stochastic hill-climbing over the value function. We posit this function as readily accessible within the BG, courtesy of top-down processing from the prefrontal areas, where goal related information is available.

Leveraging this value function, we derive a value difference signal regulating BG pathways and connections within the STN-GPe network. The value difference signal in our model acts similar to the dopamine signal that is used to switch between the direct and indirect pathways by its discriminative actions on the D1 and D2 cells of the striatum. The dopamine signal also influences STN dynamics, representing the projections of SNc to STN. In order to produce Parkinsonian tremor, we manipulate this signal, mimicking reduced dopamine levels, which profoundly influences the dynamics of the STN-GPe subsystem.

Consistent with experimental findings, our model demonstrates an increase in beta power within the STN’s local field potential (LFP) under reduced dopamine conditions [62]. Additionally, we observe that pulsatile deep brain stimulation (DBS) to STN neurons, contingent upon parameters such as amplitude, frequency, and pulse width, significantly diminishes the beta peak in the STN LFP signal, thereby mitigating tremor. The current model has the potential to be expanded to study the effect of DBS on other motor symptoms as well.

However, STN DBS is not without its side effects. Studies have reported emergence of impulsive control disorder (ICD) following the stimulation of STN [69], [70], [71], [72], [73], [74]. The connection between the STN DBS and ICD is debatable as some studies also reveal that not a large percentage of people undegoing DBS has experienced a cognitive decline [75], [76] where as some studies report a favourable effect on cognitive performance [77], [78]. However, in spite of all the diverse results it would interesting to explore the effect DBS treatment on motor and cognitive symptoms simultaneously.

In the future our aim is to expand the proposed model into a comprehensive DBS framework, encompassing all the major BG loops especially focusing on motor and cognitive symptoms. While DBS effectively attenuates motor symptoms, concerns persist regarding its impact on cognitive function. Existing studies have yet to fully elucidate the precise role of DBS within basal ganglia subcortical structures in alleviating Parkinsonian symptoms and the underlying mechanisms driving cognitive decline [79], [80].

Therefore, our next step involves refining this model to unravel the nuanced interplay between DBS, motor symptoms, and cognitive function. By meticulously regulating model parameters, we strive to develop an optimized intervention capable of addressing both motor and cognitive impairments comprehensively. Another expansion of the model is to develop a control loop that optimizes the right amount of medication and stimulation intervention at various stages of the progression of the disease. In this process there is a potential scope of expanding the current DBS model to explore different target areas for optimum therapeutical benefit. Multiple target areas commonly used for DBS intervention are ventralis intermedialis (Vim) of thalamus, Globus Pallidus Internus (GPi) and Sub Thalamic Nucleus (STN) [81], [82], [83].

