## Supplementary Data for "A Computational Model of Deep Brain Stimulation for Parkinson’s Disease Tremor"

### S1. Training of the cortico-basal ganglia model

We use a cortico-BG (CBG) model that controls a two-linked arm model in order to simulate reaching movements as shown in (Figure S1). In this section we will first introduce the CBG model, the training of the outer cortico-basal ganglia loop and the model parameters. The training scheme and the model is adapted from [1], [2], [3]

#### 1.1. Cortico-basal Ganglia Model

The CBG model used in our study comprises of an outer sensory motor loop, which interacts with the BG circuitry. The sensory motor loop comprises of an arm model, proprioceptive cortex (PC), motor cortex (MC), the prefrontal cortex (PFC) and the BG. The kinematic arm model performs reaching movements based on the activations it receives and the PC estimates the current arm position and sends the feedback to MC, which integrates this signal along with the goal information from the PFC and the error corrected signal from the BG, which does its own internal processing before sending back the corrected signal to the motor cortex. The motor cortex then sends the next motor command to the arm via the spinal motor neurons and this process continues until the arm reaches the target or the time out is reached. More details about the sensory motor loop and the arm model are discussed in the following sections.

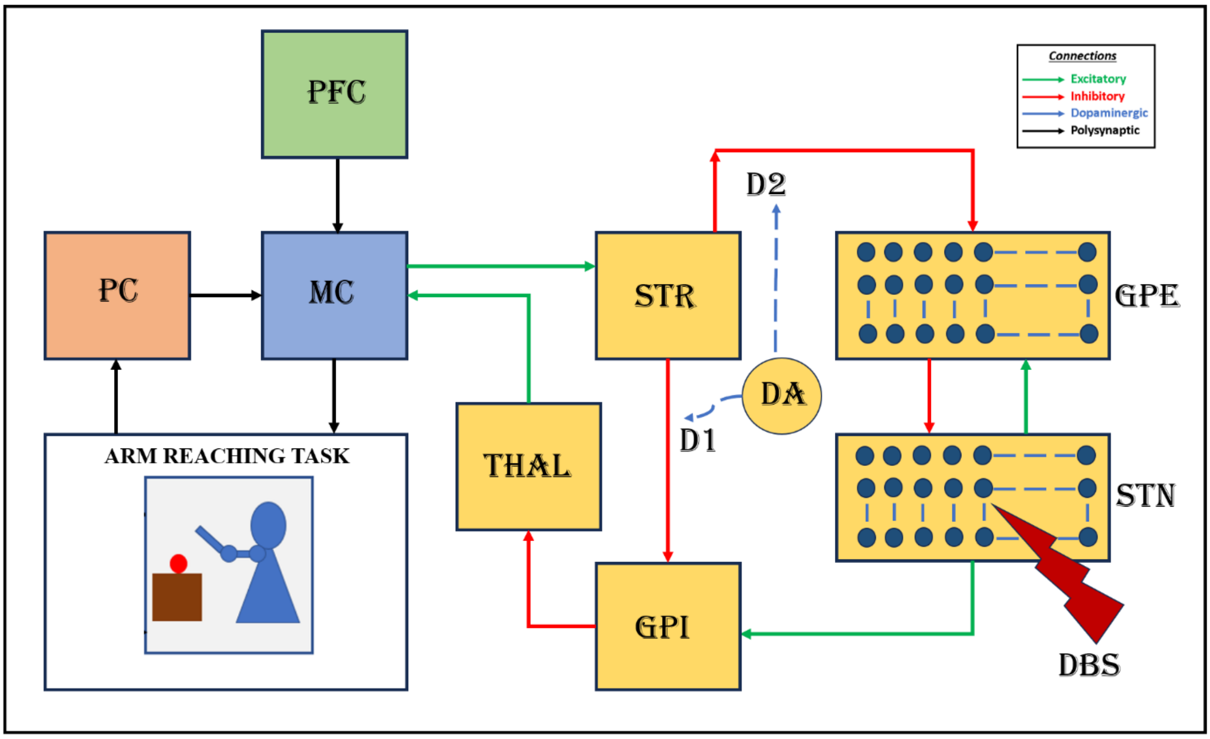

**Figure S1.** Block diagram of the proposed cortico-basal ganglia model. The model consists of a 2-link arm model, the proprioceptive cortex (PC), the prefrontal cortex (PFC), the motor cortex (MC), and Basal ganglia (BG). Here the input nucleus Striatum, the output nucleus Globus pallidus interna (GPi), the Globus pallidus externa (GPe), the subthalamic nucleus (STN), and the thalamus (THAL) constitute the BG. MC integrates the inputs received from the prefrontal cortex (PFC) and the proprioceptive cortex (PC) along with the feedback signal from BG and sends the signal to the arm via the spinal motor neurons.

##### 1.1.1. Training of the outer sensory-motor loop

The training of the sensory-motor loop comprises of the below steps.

1. First, we generate $n$ random activation vectors for the motor neurons resulting in various arm movements. This yields n different arm configurations, each providing a feature vector of muscle lengths $(M^L)$.

2. $M^L$ is given as input to the PC that undergoes learning using self-organizing maps to form a sensory activation map.
3. The PC output is the input to the MC, and MC output is given as input to MN so that the desired activation vector ($\phi^{MN}$)- initially given to the arm is obtained back thereby closing the loop.
4. In the above three steps, first the sensory activation map was formed by training the connections between the arm and PC. Then, the connections between PC and MC are trained, and in the third stage, the connections between the MC and MN are trained.
5. At the final step, BG is introduced and PFC to MC connections are trained (Figure S2).

Before discussing the details of the training mechanisms involved in the cortico-basal ganglia model used in our model, we will first define the arm model and its important parameters and then discuss how the muscle length vector is calculated.

##### 1.1.2. Obtaining the Muscle length vector for different activations

As mentioned in the previous section the training of the model starts with obtaining the muscle length vector for various activations of the motor neurons. The arm model simulates the movement of the arm in the two dimensional space [4], [5] (Figure S2). In our model we consider two joints – the shoulder and the elbow. These joints are controlled by an antagonist (An) and an agonist (Ag) muscle pair and the activations to these muscle groups ($\phi_{SE}^{MN}$) are transformed into joint angles for both shoulder and the elbow as follows,

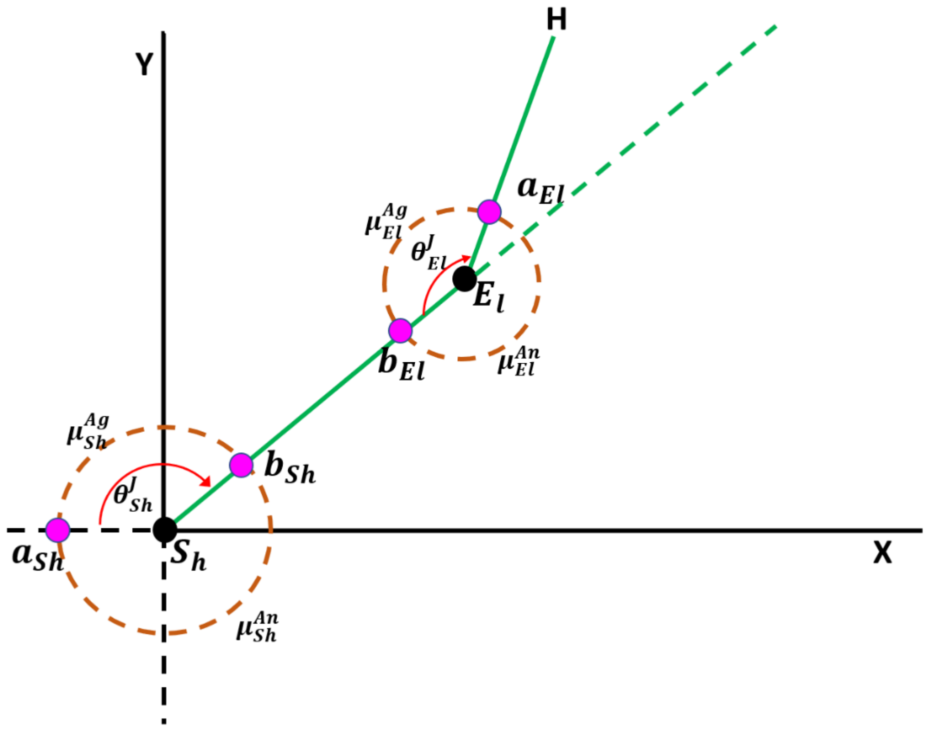

***Figure S1: The rm model.*** *$S_h$ and $E_l$ are the shoulder and elbow joints; the end effector position is marked as $H$; The axes of the plane are labeled as $X$ & $Y$; $l_e$ and $l_S$ are the distances between the (H and $E_l$) and ($E_l$ and $S_h$) respectively; $\theta_{Sh}^{J}$ and $\theta_{El}^{J}$ represents the angles at the shoulder and elbow respectively; $[\mu_{Ag}^{Sh}, \mu_{An}^{Sh}, \mu_{Ag}^{El}, \mu_{An}^{El}]$ represents the muscle length vector for agonist and antagonist pairs of shoulder and elbow muscles, respectively; $[a_{Sh}, b_{Sh}, a_{El}, b_{El}]$ represents the distance between ($S_h$ and $M_1$ or $M_2$ moment lever) and ($S_h$ and $M_3$ or $M_4$ moment lever) respectively.*

$$\theta_{Sh}^{J}(t) = \left(\phi_{Sh_{Ag}}^{MN}(t) - \phi_{Sh_{An}}^{MN}(t)\right)\frac{\pi}{2} + \frac{\pi}{2} \tag{1}$$

$$\theta_{El}^{J}(t) = \left(\phi_{El_{Ag}}^{MN}(t) - \phi_{El_{An}}^{MN}(t)\right)\frac{\pi}{2} + \frac{\pi}{2} \tag{2}$$

where, $\theta_{El}^{J}$ and $\theta_{Sh}^{J}$ are the elbow and shoulder joint angles with respect to the horizontal axis (Figure S1) and and shoulder length $(l_{Sh})$ respectively. $\phi_{S_{Ag}}^{MN}$, $\phi_{S_{An}}^{MN}$ , $\phi_{El_{Ag}}^{MN}$ and $\phi_{El_{An}}^{MN}$ are the muscle activation of the agonist and the antagonist muscles of the shoulder and elbow muscles respectively.

The joint angles defines the coverage of the arm in the two dimensional space and are used to calculate the muscle lengths as given in equations(3-6) below. This process is represented in (Figure S3)

$$\mu_{Ag}^{Sh}(t) = \sqrt{a_{Sh}^2 + b_{Sh}^2 + 2a_{Sh}b_{Sh}\cos(\theta_S^J)} \tag{3}$$

$$\mu_{An}^{Sh}(t) = \sqrt{a_{Sh}^2 + b_{Sh}^2 - 2a_{Sh}b_{Sh}\cos(\theta_{Sh}^J)} \tag{4}$$

$$\mu_{Ag}^{El}(t) = \sqrt{a_{El}^2 + b_{El}^2 + 2a_{El}b_{El}\cos(\theta_{El}^J)} \tag{5}$$

$$\mu_{AN}^{El}(t) = \sqrt{a_{El}^2 + b_{El}^2 - 2a_{El}b_{El}\cos(\theta_{El}^J)} \tag{6}$$

where, $\mu_{Ag}^{Sh}$, $\mu_{An}^{Sh}$, $\mu_{Ag}^{El}$, and $\mu_{An}^{El}$ are the agonist and antagonist muscle lengths of shoulder and elbow, respectively, $a_{Sh}$ and $b_{Sh}$ are the distance between shoulder joint center and $M_1$ or $M_2$ moment lever, and, $a_{El}$ and $b_{El}$ are the distance between elbow joint center and $M_3$ or $M_4$ moment lever respectively.

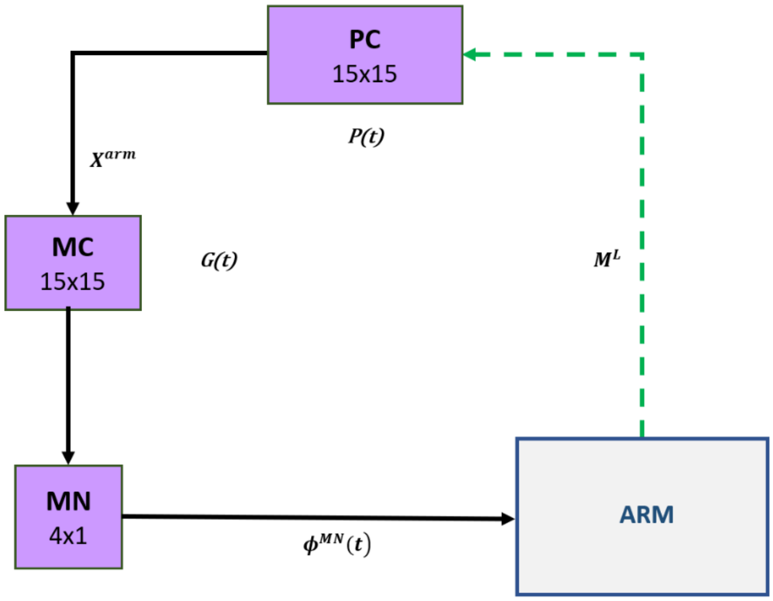

***Figure S3: The training schema for ARM-PC connections (Sensory-motor loop).*** *In the figure, the dashed arrows indicate the connections that are being trained. PC, proprioceptive cortex; MC, motor cortex; MN, motor neuron; PFC, prefrontal cortex; BG, basal ganglia;* $X^{arm}$*, the current arm position;* $\phi^{MN}$*, the motor neuron activations;* $M^L$*, muscle lengths;* $P(t)$*, the PC output;* $\Delta G(t)$*, the MC output.*

Using these muscle lengths in the form of a four-dimensional vector $\left(M^L = [\mu_{Ag}^{Sh}\ \mu_{An}^{Sh}\ \mu_{Ag}^{El}\ \mu_{An}^{El}]\right)$, a sensory (proprioceptive) map of the arm was generated. The end effector position of the arm $(X^{arm} = [x_1^{arm}\ x_2^{arm}])$ in the two-dimensional space is calculated as shown in equations(7-8) ,

$$x_1^{arm} = (l_{Sh} - a_{Sh})\cos(\theta_{Sh}^J) + l_{El}\cos(\theta_{Sh}^J + \theta_{El}^J) \quad (7)$$

$$x_2^{arm} = (l_{Sh} - a_{Sh})\sin(\theta_{Sh}^J) + l_{El}\sin(\theta_{Sh}^J + \theta_{El}^J) \quad (8)$$

$$M^L = [\mu_{Ag}^{Sh}\ \mu_{An}^{Sh}\ \mu_{Ag}^{El}\ \mu_{An}^{El}] \quad (9)$$

where, $\theta_{Sh}^J$ and $\theta_{El}^J$ are the joint angles of shoulder and elbow with respect to the horizontal axis (Figure S1) and shoulder length $(l_{Sh})$, respectively in two-dimensional space, $l_{Sh}$ is the distance between the shoulder joint center $(S)$ and elbow joint center $(E)$, $l_{El}$ is the distance between the elbow joint center $(E)$ and end effector $(H)$. $M^L$ is the muscle length vector.

In order to train the PC to MC connections as shown in (Figure S4), we need to first obtain the sensory activation map and the activation map for the PFC. These two processes are discussed in subsections (1.1.3 and 1.1.4) below.

##### 1.1.3. Obtaining the sensory activation maps

Using the $M^L$ computed above, we proceed with obtaining the sensory activation map. PC receives the input from the arm model and it sends the feedback signal to the MC. It is modelled as a self-organizing map (SOM) [6] where sensory map of the arm is created. Below equation (10) shows the activity of the single neuron of the PC,

$$U_{ij}^{PC}(t) = exp\left(\frac{-\left\|M^L(t) - W_{ij}^{PC}\right\|^2}{(\sigma^{PC}).(\sigma^{PC})}\right) \quad (10)$$

where, $W_{ij}^{PC}$ is the weight between $M^L$ and $(i,j)^{th}$ neuron of the two-dimensional PC network, $M^L$ is the muscle length vector and the spatial distance within which the response of the PC SOM shows sensitivity is given by $\sigma^{PC}$.

##### 1.1.4. Training of the Prefrontal Cortex (PFC)

PFC provides the goal position. Biologically the goal position is obtained by the visual feedback, where as in our model we fix the goal position as $X^{targ}$. The initial signal from the motor cortex (MC) to motor neurons (MN) is initially facilitated by the PFC and gradually with learning the BG circuitry takes over. All possible locations where the arm has the possibility to reach constitutes the features of PFC. The PFC is trained using the target vector and the activity of neuron at $(i, j)^{th}$ location in the PFC is given as in equations(11).

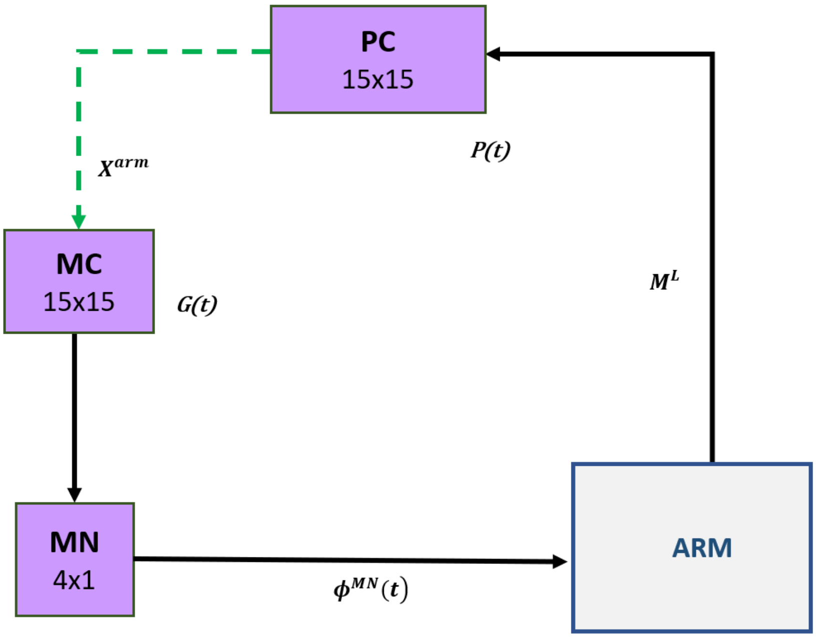

***Figure S3: The training schema for PC-MC connections (Sensory-motor loop).*** *In the figure, the dashed arrows indicate the connections that are being trained. PC, proprioceptive cortex; MC, motor cortex; MN, motor neuron; PFC, prefrontal cortex; BG, basal ganglia;* $X^{arm}$*, the current arm position;* $\phi^{MN}$*, the motor neuron activations;* $M_L$*, muscle lengths;* $P(t)$*, the PC output;* $\Delta G(t)$*, the MC output.*

$$U_{ij}^{PFC}(t) = exp\left(\frac{-\left\|X^{targ}(t) - W_{ij}^{PFC}\right\|^2}{(\sigma^{PFC}).(\sigma^{PFC})}\right) \tag{11}$$

$W_{ij}^{PFC}$ is the weight between the target position vector and $(i, j)^{th}$ neuron and the 2-D PFC network and $\sigma^{PFC}$ is the width of the PFC SOM response.

##### 1.1.5. Training MC to Motor neuron (MN) connection

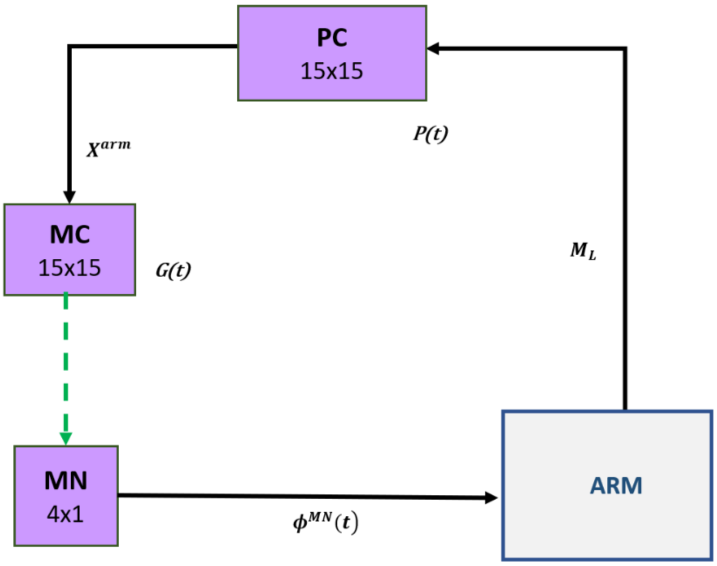

***Figure S5: The training schema for MC-MN connections (Sensory-motor loop).*** *In the figure, the dashed arrows indicate the connections that are being trained. PC, proprioceptive cortex; MC, motor cortex; MN, motor neuron; PFC, prefrontal cortex; BG, basal ganglia;* $X^{arm}$*, the current arm position;* $\phi^{MN}$*, the motor neuron activations;* $M_L$*, muscle lengths;* $P(t)$*, the PC output;* $\Delta G(t)$*, the MC output.*

The MC output projects to the four MN neurons, which drives the muscle activations in order to make the arm movements. The activation of the MN layer is given by equation (12),

$$\phi^{MN} = A^{MN} W^{MC\rightarrow MN} G^{MC}(t) \tag{12}$$

where, $A^{MN}$ is the gain of MN, $W^{MC\rightarrow MN}$ is the weight matrix between MC and MN layer, and $G^{MC}$ is the output activity of MC CANN.

To close the sensory-motor loop, we perform a comparison with the initial activation to the MN layer that was used to produce desired activation $\phi_D^{MN}(t)$. The weights between the MN and MC are trained in a supervised manner by comparing the network derived MN activation $\phi^{MN}(t)$ to the desired activation $\phi_D^{MN}(t)$. This gives a loop which is consistent in mapping the external arm space to the neuronal space and vice. The connection between MC and MN is trained as represented in (Figure S5) and is given by equation (13),

$$\Delta W^{MC\rightarrow MN} = \eta^{MC\rightarrow MN}\left(\phi_D^{MN}(t) - \phi^{MN}(t)\right)G^{MC}(t) \tag{13}$$

where, $\eta^{MC\rightarrow MN}$ is the learning rate between MC and MN, $\phi_D^{MN}$ is the desired MN activation required for the arm to reach the target and $\phi^{MN}$ is the network-derived MN activation due to

MC, and $G_{MC}$ is the output activity of MC. The training schema for the outer loop (sensory-motor loop) is described in section S3 of the supplementary information.

##### 1.1.6. Motor Cortex (MC) and MC to PFC training

The MC receives inputs from the PFC, the PC and the BG. MC is modeled as a combination of self organizing map and continuous attractor neural network (CANN) [7]. The total input received at the MC, $I^{MC}$, is contributed by the PC, the PFC and the BG as shown in equation (14) below.

$$I^{MC} = A^{PFC}.G^{PFC} + A^{PC}.G^{PC} + A^{BG}.G^{BG} \tag{14}$$

where, ($A^{PFC}$, $A^{PC}$, $A^{BG}$ ) and ($G^{PFC}$, $G^{PC}$, $G^{BG}$) represents the gains and output activities of PFC, PC and BG respectively. The PC activity used to generate feature maps in MC is given below where the activation of the $(i,j)^{th}$node in the SOM part of the MC $\left(G_{PC,ij}\right)$ is given as in equation (15) below,

$$G_{PC,ij}(t) = exp\left(\frac{-\left\|U^{PC}(t) - W_{ij}^{MC}\right\|^2}{(\sigma^{MC}).(\sigma^{MC})}\right) \tag{15}$$

where, $U^{PC}$ is the output activity of PC SOM network, $W_{ij}^{MC}$ is the weight between the PC network and $(i,j)^{th}$ neuron of the two-dimensional MC network, and the spatial distance within which the response of the MC SOM shows sensitivity is given by $\sigma^{MC}$.

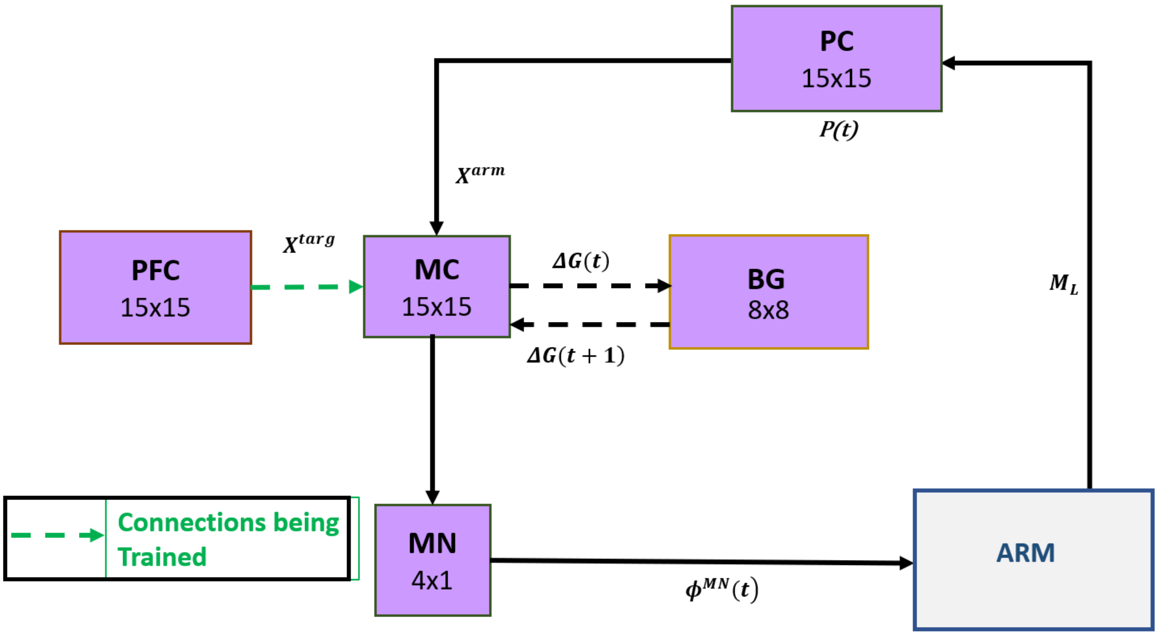

***Figure S6: The training schema for PFC to MC connections (sensory-motor loop.*** *BG module is introduced and the PFC to MC connections are trained. The dashed arrows indicate the connections that are being trained. PC, proprioceptive cortex; MC, motor cortex; MN, motor neuron; PFC, prefrontal cortex; BG, basal ganglia;* $X^{targ}$*, the target position;* $X^{arm}$*, the current arm position;* $\phi^{MN}$*, the motor neuron activations;* $M_L$*, muscle lengths;* $P(t)$*, the PC output;* $\Delta G(t)$*, the MC output;* $\Delta G(t+1)$*, the BG-derived activity of thalamus.*

The input from PFC to MC ($G_{PFC}$) is the product of weight matrix ($W^{PFC\to MC}$) and the output activity of PFC SOM is given by equation (16) below,

$$G_{PFC}(t) = W^{PFC\to MC} * U^{PFC}(t) \tag{16}$$

where, $U^{PFC}$ is the output activity of PFC SOM network, $W^{PFC\to MC}$ is the weight matrix between PFC and MC.

The output of MC is defined by equation (17) below,

$$G^{MC}(t) = \frac{\left(g^{MC}\right)^2}{1+\left(\frac{2\pi}{\left(N^{MC}\right)^2}\right)b^{MC}\sum\left(g^{MC}\right)^2} \tag{17}$$

where, $N^{MC}$ defines the size of the network, $b^{MC}$ is a constant and $g^{MC}$ represents the intrinsic dynamics of the MC neurons and it is as given in the equation (18) below,

$$\tau_{MC}\frac{dg^{MC}}{dt} = -g^{MC} + W_{MC}^{C} \otimes G^{MC} + I^{MC} \tag{18}$$

where, the weight kernel $W_{MC}^{C}$ represents the lateral connectivity among the MC neurons as given in equation (19), whose dynamics are defined by local excitation and global inhibition, and ⊗ represents the convolutional operation.

$$W_{MC,i,j}^{C} = A_{lat}^{C} \exp\left(\frac{-\|(i^{MC} - i^{h}) + (j^{MC} - j^{h})\|^{2}}{2(\sigma_{lat}^{C})^{2}}\right) - K^{C} \tag{19}$$

where, $[i^{MC}, j^{MC}]$ are the position of the neurons in MC, $[i^{h}, j^{h}]$ is the position of the neuron in the centre, $A_{lat}^{C}$ is the strength of the excitatory connections, $K^{C}$ is the global inhibition constant and $\sigma_{lat}^{C}$ is the radius of the excitatory connections.

When the arm reaches the target ($\epsilon < 0.1$), the connections between the PFC and MC are trained and is as given in equation (20) below,

$$\Delta W_{PFC\rightarrow MC} = \eta_{PFC\rightarrow MC}\left(G_{targ}^{MC}(t) - G_{PFC}^{MC}(t)\right)U^{PFC}(t) \tag{20}$$

where, $\eta_{PFC\rightarrow MC}$ is the learning rate between PFC and MC, $G_{targ}^{MC}$ is the MC activation required for the arm to reach the target, and $G_{PFC}^{MC}(G_{PFC})$ is the MC activation due to PFC.

#### 2 The Model parameters

The model parameters used in our model is as given in Table S1 below. The parameters [$a^{gpe}$, $b^{gpe}$, $c^{gpe}$, $d^{gpe}$] and [$a^{STN}$, $b^{STN}$, $c^{STN}$, $d^{STN}$] represents the Izhikevich parameters using which the spiking neurons were modeled. $I_{STN}$ and $I_{GPe}$ are the STN and GPe currents. $E_{AMPA/NMDA/GABA}$ are the resting membrane potentials of different receptors associated with STN and GPe. $\tau_{AMPA/NMDA/GABA}$ are the time decay constants of the receptors. $W_{STN}^{max}$ and $W_{GPe}^{max}$ are the maximum collateral synaptic strength. DA is the dopamine signal, $A^{D1} and\ A^{D2}$ are the synaptic weight between D1 striatum and GPi and synaptic weight between D2 Striatum and GPe respectively.

**Table 1. Model parameters**

| **Parameter** | **Values with description** |
|---|---|

| | | |
|---|---|---|
| | STN | GPe |
| Izhikevich parameters | $a^{gpe}$=0.005, $b^{gpe}$=0.265, $c^{gpe}$=-65, $d^{gpe}$=1.5 | $a^{STN}$=0.1, $b^{STN}$=0.2, $c^{STN}$=-65, $d^{STN}$=2 |
| External current (I) | $I_{STN}$ = 20 pA | $I_{GPe}$ = 4.5 pA |
| $A^{D1}$ | 7.5 | Synaptic weight between D1 striatum and GPi |
| $A^{D2}$ | 2 (HC and Tremor); (<0.4) Rigidity | Synaptic weight between D2 striatum and GPe |
| $A^{GPe}$ | 1 | |
| DA | 0.1 to 0.9 in increments of 0.1 | |
| $Mg^{2+}$ | 1nM | Concentration of Magnesium ions in nM |
| $E_{AMPA}$ | 0mv | Synaptic potential of AMPA receptor-associated channel |
| $E_{NMDA}$ | 0mV | Synaptic potential of NMDA receptor-associated channel |
| $E_{GABA}$ | -60mV | Synaptic potential of GABA receptor-associated channel |
| $\tau_{AMPA}$ | 6ms | Time decay constant for AMPA receptor |
| $\tau_{NMDA}$ | 160ms | Time decay constant for NMDA receptor |
| $\tau_{GABA}$ | 4ms | Time decay constant for GABA receptor |
| $W_{GPe}^{max}$ | 1 | Synaptic strength within GPe laterals |
| $W_{STN}^{max}$ | 0.51 | Synaptic strength within STN laterals |

#### S2. Synchrony

The synchronization measure, $R^{syn}$, is calculated using the instantaneous phases of the neurons in the subpopulation of STN and GPe neurons. $R^{syn}$ gives a measure of the correlation among the neurons. In healthy state (HC), the synchrony will be less, while in the disease state the synchrony will be high. Park et al (2011) report the presence of intermittent synchrony between STN neurons and its Local field potentials (LFP), recorded using multiunit activity electrodes from PD patients undergoing DBS surgery (Park, Worth et al. 2011). They also calculated the duration of synchronized and desynchronized events in neuronal activity by estimating transition rates, which were obtained with the help of first return maps plotted using phase of neurons [8], [9] . To observe how dopamine changes synchrony in STN-GPe, we calculated the phases of individual neurons as defined in [10] .

The phase of $j^{th}$ neuron was calculated as follows,

$$\phi_j(t) = 2.\pi.\frac{(T_{j,k}-t_{j,k})}{(t_{j,k+1}-t_{j,k})} \tag{21}$$

$$R^{syn}(t).e^{i\theta(t)} = (1/N)\sum_{j=1}^{N} e^{i\phi_j(t)}) \tag{22}$$

where, $t_{j,\,k}$ and $t_{j,\,k+1}$ are the onset times of $k^{th}$ and $k+1^{th}$ spike of the $j^{th}$ neuron $T_{j,k} = [t_{j,k}, t_{j,k+1}]$, $\phi_j(t)$= Phase of $j^{th}$ neuron at time'$t$', $R^{sync}$ is the synchronization measure $0 \leq R^{sync} \leq 1$ , $\theta$=Average phase of neurons, N=total number of neurons in the network.

$t_{j,k}$
